# Cell types and molecular architecture of the octopus visual system

**DOI:** 10.1101/2022.06.11.495763

**Authors:** Jeremea O. Songco-Casey, Gabrielle C. Coffing, Denise M. Piscopo, Judit R. Pungor, Andrew D. Kern, Adam C. Miller, Cristopher M. Niell

## Abstract

Cephalopods have a remarkable visual system, with a camera-type eye, high acuity vision, and a wide range of sophisticated visual behaviors. However, the cephalopod brain is organized dramatically differently from that of vertebrates, as well as other invertebrates, and little is known regarding the cell types and molecular determinants of their visual system organization beyond neuroanatomical descriptions. Here we present a comprehensive single-cell molecular atlas of the octopus optic lobe, which is the primary visual processing structure in the cephalopod brain. We combined single-cell RNA sequencing with RNA fluorescence in situ hybridization to both identify putative molecular cell types and determine their anatomical and spatial organization within the optic lobe. Our results reveal six major neuronal cell classes identified by neurotransmitter/neuropeptide usage, in addition to non-neuronal and immature neuronal populations. Moreover, we find that additional markers divide these neuronal classes into subtypes with distinct anatomical localizations, revealing cell type diversity and a detailed laminar organization within the optic lobe. We also delineate the immature neurons within this continuously growing tissue into subtypes defined by evolutionarily conserved fate specification genes as well as novel cephalopod- and octopus-specific genes. Together, these findings outline the organizational logic of the octopus visual system, based on functional determinants, laminar identity, and developmental markers/pathways. The resulting atlas presented here delineates the “parts list” of the neural circuits used for vision in octopus, providing a platform for investigations into the development and function of the octopus visual system as well as the evolution of visual processing.

**Highlights:**

- Single-cell RNA sequencing coupled with RNA fluorescence in situ hybridization produces a molecular taxonomy of cell types in the octopus visual system.
- Six major neuronal cell classes are delineated based on neurotransmitters/neuropeptides, and are further subdivided based on laminar organization and additional marker genes.
- Immature neurons are divided into multiple transcriptional subgroups that correspond to mature cell types, delineated by expression of genes known for their developmental roles in other organisms as well as apparent novel genes.
- This atlas provides the foundation for future studies of the function, development, and comparative evolution of visual processing in cephalopods.

## Introduction

Cephalopods represent a unique branch of the animal kingdom for the study of visual processing. Coleoid cephalopods (octopuses, squids, and cuttlefish) have the largest brain among invertebrates^1–3^, with two-thirds of their central brain dedicated to visual processing^4,5^. Their visual system facilitates a range of behaviors such as navigation, prey capture, and complex camouflage^1,6–8^. However, the neural basis of central visual processing in cephalopods is largely unknown.

Despite diverging over 500 million years ago, octopuses independently evolved camera-type eyes similar to those of vertebrates^9^, with a pupil and a lens that focuses light to form an image on the retina. While this is often viewed as a “textbook” example of convergent evolution^10,11^, the neural organization of structures that process visual information is dramatically different in the two lineages. In contrast to the vertebrate retina, which is an intricate neural circuit that performs complex computations through diverse cell types, the octopus retina consists only of photoreceptors, with a small population of cells that stratify horizontally, traditionally referred to as supporting cells^12,13^. The photoreceptors themselves send axons into the central brain, targeting the optic lobes that lie directly behind the eyes^12^ (Fig. 1A). The optic lobes, which make up approximately two-thirds of the octopus’ central brain^14,15^, are where the bulk of visual processing is hypothesized to occur^14–16^, and contain outputs projecting to deeper brain regions involved in learning/memory and motor behavior^14,17–21^. Thus, while morphological similarities exist between the vertebrate and cephalopod camera-type eyes, there are clear differences in large-scale retinal and brain organization between the two systems. On the other hand, the octopus optic lobe is a layered structure (Fig. 1B-C), suggesting that the optic lobes may represent, at the cellular level, analogous neural circuitry to that which resides in the layered vertebrate retina^3^.

**Figure 1.**
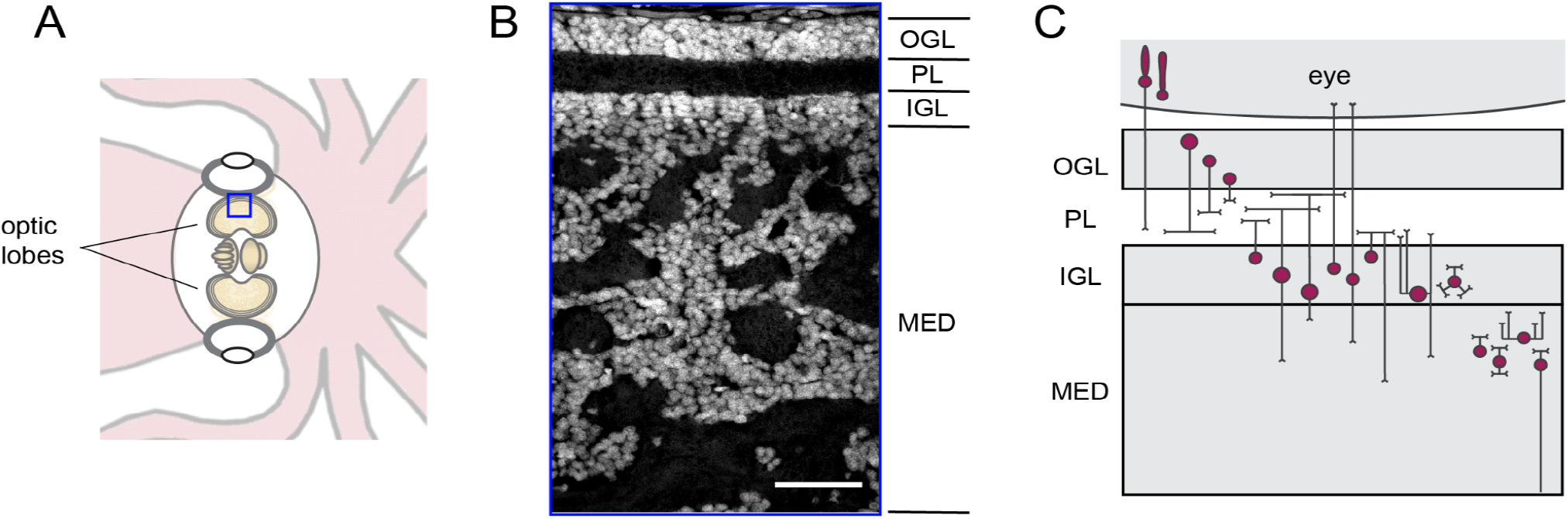
Laminar organization of *Octopus bimaculoides* optic lobes. (A) Schematic showing organization of the visual system including the eye and optic lobes. Blue box denotes the region shown in B. (B) Laminar organization of the optic lobe demonstrated by nuclear staining (DAPI) of a cross-section. OGL, outer granular layer; PL, plexiform layer; IGL, inner granular layer; MED, medulla. Scale bar indicates 50 um. (C) Schematic of the anatomical organization of the eye and optic lobe, in terms of neuronal morphologies, adapted from ^5^.

Histological studies using Golgi stains^5^ have provided a description of the optic lobe’s anatomical organization and neuronal morphologies (Fig. 1C), which we briefly summarize here. The outermost region of the optic lobe is a cell body layer, termed the outer granular layer (OGL). The OGL contains cells which have multipolar processes without clear differentiation between axons and dendrites, traditionally referred to as amacrine cells based on their morphology^20,22^. These amacrine cells ramify within the plexiform layer below the OGL. The plexiform layer is a dense neuropil and is the primary termination of photoreceptor axons, optic lobe neuronal processes, and projections from deeper brain regions^14,23^. Below the plexiform layer is another cell body layer, the inner granular layer (IGL), which has a varied population of neurons including (1) a large number of neurons with bipolar morphology restrained within their optical column, with dendrites in the plexiform layer and axons that project further within the optic lobe, (2) a set of neurons that send centrifugal axons back to the retina, and (3) another group of cells with amacrine morphology with processes in the plexiform layer^14,20^. Finally, a deeper structure termed the medulla comprises the bulk of the optic lobe. The cells within the medulla are organized in a branching tree-like fashion which, when viewed in a cross-section, appears as islands of cell bodies surrounded by neuropil^24^. The medulla contains both unipolar and multipolar neurons, some of which project axons out of the optic lobe to central brain structures. The superficial region of the medulla contains cell bodies organized into columns and is traditionally referred to as the outer radial columnar zone, whereas the deeper region of the medulla includes processes that extend tangentially and is therefore termed the central tangential zone^5^. Based on this anatomical organization, it has been hypothesized that the outer layers of the optic lobe (OGL, plexiform, IGL) may perform similar functions to that of the vertebrate inner retina, leading it to be termed the “deep retina”. Meanwhile the medulla may engage in higher order processing analogous to central visual areas in other species^14^.

While these anatomical classes of the cells in the octopus optic lobe suggest an organizational foundation, a full understanding of the neural circuitry necessitates knowledge of the molecular identities of the underlying cell types. These molecular signatures include both functional determinants, such as neurotransmitter and receptor repertoires, and developmental determinants, such as transcription factors and adhesion molecules that establish cell type identity and connectivity. Recent molecular taxonomies in other species have provided new insight into the circuit organization of a number of brain regions including the fly visual system^25^, the mouse and primate retina^26,27^ and the mouse visual cortex^28^. These studies have further revealed extensive diversity even within previously studied cell types. We therefore sought to create a systematic molecular characterization of the optic lobe cell types in the octopus visual system.

Here we combine single-cell RNA sequencing (scRNA-seq) of the octopus optic lobe with multiplexed RNA fluorescence in situ hybridization (FISH) to determine the molecular cell types of the octopus optic lobe and their spatial organization, resulting in a comprehensive atlas of the octopus visual system. This work reveals a wide range of diverse cell types with intricate laminar and graded organization, intersected with potential functional repertoires (neurotransmitters/neuropeptides), and uncovers gene expression information for putative subtypes of immature neuronal populations. This atlas provides the framework for understanding visual processing in cephalopods from the perspectives of functional circuits, developmental logic, and comparison of cell types across evolution.

## Results

### Single-cell RNA sequencing of the octopus optic lobe

We performed scRNA-seq in juvenile (1.5 months of age) *Octopus bimaculoides*. Despite their body and brain continuing to grow throughout their lifetime (1-2 years), the overall body organization, behavior, and visual system of *O. bimaculoides* are mature by this age^29,30^, allowing us to identify neural circuitry involved in a fully functioning, yet still growing, visual system. In order to determine the molecularly defined cell types of the octopus visual system, we dissected the optic lobes from two individuals and dissociated the tissue into single cell suspensions, with each animal processed separately as a biological replicate. Following dissociation, we sequenced cells using Chromium 10x and used Cell Ranger^31^ to align reads to an updated genome assembly and gene annotations (see Methods, Fig. S1 and Table S1). These updated gene annotations were necessary to resolve limitations in the original genome for more accurate single-cell mapping^32^ (Fig. S1). We then applied standard filtering, normalization, clustering, and integration (see Methods) to process single cell gene expression using Seurat^31,33^, identifying a total of 28,855 cells. We performed dimensionality reduction with principal components analysis, followed by k-nearest neighbors clustering, which resulted in a total of 41 clusters. We found that each biological replicate contributed cells to all of the identified clusters with similar proportions, and that the quality of the data was similar between the samples, supporting the reproducibility of the approach (Fig. S2).

We first sought to broadly characterize the identified clusters in terms of neuronal and non-neuronal populations (Fig. S3). We used a homologous sequence identifier, OrthoFinder^34^, to assign gene-family and orthology relationships of our updated *O. bimaculoides* gene annotations compared to *Drosophila*, vertebrates, and other cephalopod species (see Methods, and Fig. S3A for example gene trees). Here, and in all figures, wherever confident orthology was found, we name the octopus genes by their gene-family identity: e.g. synaptotagmin (*syt*), as summarized in Table S2.

We expected a large proportion of the cells to have high expression of genes related to mature neurons as well as genes related to glia, vasculature, and likely developing neurons, given that the octopus brain continues to grow and add neurons throughout its lifetime^35^. To identify neurons within our scRNA-seq data, we first looked at the expression of genes related to synaptic vesicle release (SNARE complex proteins and *syt*). We reasoned that these evolutionarily-conserved gene families contained markers that are expressed in mature neurons across other lineages and were likely to be expressed in octopus neurons as well. We found that most clusters expressed the *O. bimaculoides* SNARE genes, suggesting that they represent neurons, while only five were likely to be non-neuronal, based on the absence of expression of these markers (Fig. S3). These non-neuronal clusters represented ∼8% of the cells and had relatively high expression of genes falling within gene families with evolutionarily conserved functions consistent with non-neuronal cell types, including proliferating cells (proliferating cell nuclear antigen; *pcna*)^36^, blood cells (acetylcholinesterase; *ache*)^37^, endothelial cells (basement membrane-specific heparan sulfate proteoglycan core protein; *hspg, collagen*)^38^, and glia (glutamine synthetase; *glul*)^39^ (Fig. S3). To map these putative non-neuronal cell transcriptional clusters to their anatomical location, we used FISH to localize the expression of marker genes for each (Fig. S3). We found several of these markers to be primarily expressed outside of the optic lobe, within the region presumed to be the white body, which is involved in hematopoiesis^40^. One prominent marker gene from these clusters that is expressed in the optic lobe is *glul*, a glial marker, which identified a population of cell bodies at the boundary of the plexiform layer and IGL, with expression extending into the neuropils of the medulla, consistent with glial organization in other invertebrate nervous systems^41^. However, since these five clusters represent non-neuronal cells, we removed them from subsequent analyses that aimed to systematically delineate the neuronal cell types. The remaining neural dataset included 26,092 cells in 33 clusters, and was the focus of further cell type analysis (Fig. 2A).

**Figure 2.**
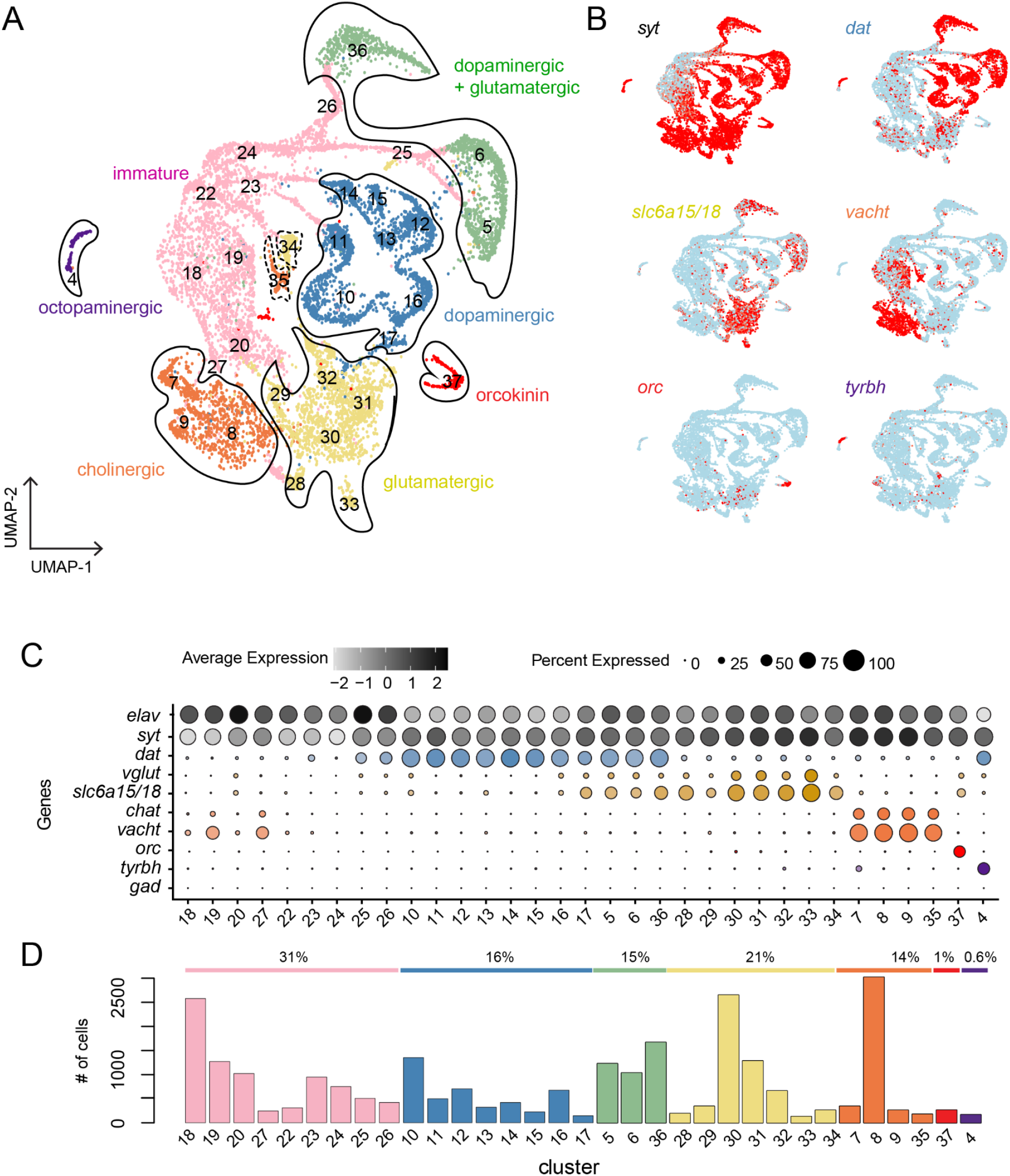
Single-cell RNA sequencing of *Octopus bimaculoides* optic lobes reveals six major neuronal classes. (A) UMAP of putative neuronal clusters. A total of 33 clusters are color-coded based on major cell classes. Each dot represents a cell and dis/similarity of transcriptional profiles is represented via distance between cells. (B) Feature plots showing expression patterns of marker genes for neurons (*syt*) and neurotransmitter phenotypes (dopaminergic cells (*dat*), glutamatergic cells (*slc6a15/18*), cholinergic cells (*vacht*), orcokinin cells (*orc*), and octopaminergic cells (*tyrbh*). (C) Dot plot of markers delineating molecular cell classes. The expression of each gene is denoted in color intensity, and the percent of cells within a cluster expressing a given gene are represented through dot size. (D) Bar graph indicating total number of cells in each cluster as colored in (A) as well as the relative proportion of each cell class across the entire population of neurons in the optic lobe.

To further explore the potential neural cell types suggested by the scRNA-seq clusters, we first examined the expression of markers for neurotransmitter usage (Fig. 2B-C). Previous work identified dopamine, glutamate, and acetylcholine as the primary neurotransmitters used in the cephalopod optic lobe^42^, and we identified orthologs corresponding to standard markers for these neurotransmitter types based on the biosynthetic and vesicular packaging pathways required for each (dopamine transporter (*dat*), vesicular glutamate transporter (*vglut*), vesicular acetylcholine transporter (*vacht*), and choline acetyltransferase (*chat*)) (Fig. 2B-C). In addition, we identified a gene in the solute carrier gene superfamily (*slc6a15/18)*, which acts to support glutamate synthesis via transport of glutamate precursors^43^, and was closely co-expressed with *vglut* (Fig. 2C). We used this as an additional marker of glutamatergic neurons, since we found a relatively low number of scRNA-seq reads aligned to *vglut* despite strong FISH signal (data not shown).

Together, expression of these neurotransmitter markers delineated the vast majority of putative neurons into four broader classes defined by either unique or combinatorial expression of these genes (Fig. 2B-C). Each of these broad categories consisted of a number of unique clusters (Fig. 2A,C), suggesting potential further transcriptional heterogeneity of cell type or state within each, which will be explored below. In addition, two smaller neuronal clusters were identified, one of which did not express markers for any neurotransmitters, but did express the conserved neuropeptide orcokinin (*orc*) (Fig. 2B) and another that expressed a combination of *dat* and a marker for octopamine synthesis, tyramine beta-hydroxylase, *tyrbh*, previously identified in octopus optic lobe neurons^44^. We did not find a significant population of GABAergic neurons (as identified by expression of glutamate decarboxylase (*gad*), Fig. 2C), consistent with previous findings of the minimal role of this neurotransmitter in the optic lobe^45^.

Finally, a large fraction of cells appear to be immature neurons, based on higher expression levels of early neural specification genes (i.e. embryonic lethal abnormal vision (*elav*) and CUG triplet repeat RNA binding protein (*cugbp*)), lower levels of genes involved in synaptic transmission such as *syt* and SNAREs, and no expression of any neurotransmitter markers (Fig. 2C, S3). As described below, these cells also expressed a diversity of evolutionarily conserved developmental genes, and distinct subgroups have expression profiles suggestive of a relationship to distinct mature cell clusters. Taken together, these findings support the idea that the scRNA-seq data captured expression profiles of unique classes of mature and immature neurons from the optic lobe. Below we use these data to delineate molecular cell types and assign them to an anatomical organization *in vivo* in the octopus optic lobe.

### A molecular and spatial taxonomy of mature neural cell types

Together, these scRNA-seq data suggest that neurotransmitter usage divides the majority of octopus optic lobe cells into four large populations – dopaminergic, co-expressing dopaminergic+glutamatergic, glutamatergic, and cholinergic neurons – along with two smaller populations expressing orcokinin and octopamine. To examine the localization of the four major neurotransmitter cell types within the anatomy of the optic lobe, we performed FISH for *dat, slc6a15/18*, and *vacht* (Fig. 3). Each of the neurotransmitter markers showed a distinct pattern of expression within the cell-body layers of the OGL, IGL, and medulla (Fig. 3B). Dopaminergic cells, identified by *dat* expression, were found within cells present in the OGL and IGL, with sparser expression in the medulla. *slc6a15/18*, a marker of the glutamatergic cell cluster, was expressed within cell bodies spread across the optic lobe, and was co-expressed with *dat* within the OGL and IGL. Finally, *vacht*, a cholinergic cell marker, was not expressed in the OGL but within a large fraction of cell bodies in the IGL and medulla. As suggested by the scRNA-seq data (Fig. 3A), glutamatergic and cholinergic neurons are segregated in their spatial expression patterns– both are in non-overlapping cells in the IGL and medulla, with *slc6a15/18+* expressed more strongly in the deeper central tangential region of the medulla and *vacht*+ expressed more strongly in the more superficial radial columnar region (Fig. 3B). Together, these mappings confirm that the scRNA-seq data identified distinct populations of neurons that correlate with distinct anatomical spatial expression patterns.

**Figure 3.**
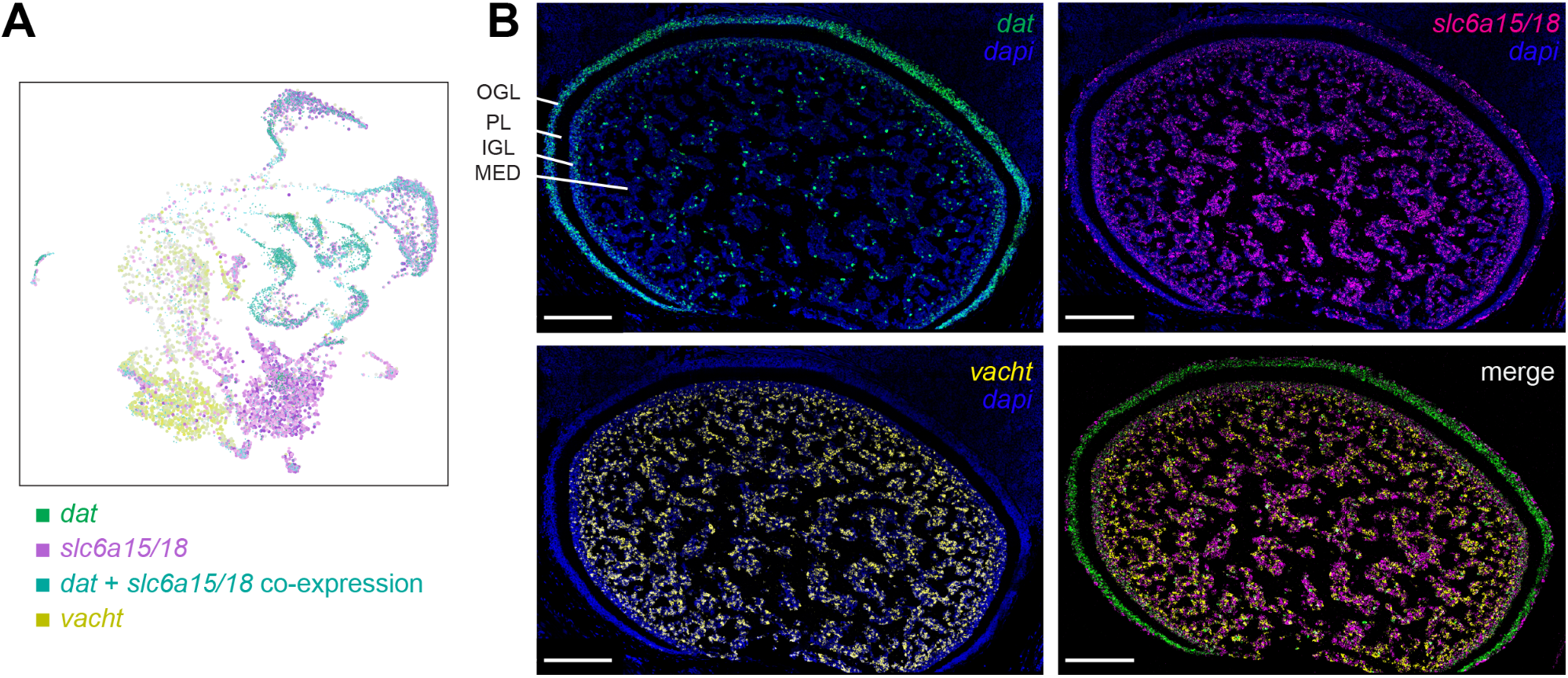
Neurotransmitter usage divides the majority of cells into four large populations. (A) UMAP of overlay of *dat, slc6a15/18*, and *vacht* expression. co-expression of *dat* and *slc6a15/18* is shown in cyan. (B) FISH of *dat, slc6a15/18, vacht*, with nuclei stained in DAPI, and merged FISH of the three neurotransmitter-related genes. Scale bars represent 100um. Here and below, merged images are shown without DAPI to emphasize gene co-expression..

Within each of the large, neurotransmitter-defined populations, further transcriptional heterogeneity was present suggesting that the clusters within each of these populations may correspond to neuronal subtypes (Fig. 2). We sought to determine if such cell types, as identified by their unique gene expression profiles, would occupy distinct anatomical locations within those of each of the broad neurotransmitter classes.

#### Dopaminergic neurons

In the scRNA-seq data, dopaminergic neurons spanned seven clusters (Fig. 2A-C), and *dat*-only neurons were predominantly localized to the OGL (Fig 3B). In order to identify subtypes within this population, we examined gene expression across these clusters and found two large subgroups defined by the exclusive expression of either the homeobox transcription factor *six3/4/5* (clusters 12-17) or the neuropeptide *fmrf1* (clusters 10-11) (Fig. 4A). FISH revealed that within the *dat+* cells in the OGL, the *six3/4/5+* population corresponds to cell bodies within a broad sub-layer of neurons in the middle of the OGL, while *fmrf1* expression corresponds to a smaller sub-layer of neurons deeper in the OGL, along the border of the plexiform layer. Thus, the *dat*-only expressing cells contain two broad groups that are differentiated by expression of a homeobox transcription factor. Notably, these are mainly localized within the OGL and hence likely represent a subset of amacrine neurons, based on Young’s anatomical study^5^.

**Figure 4.**
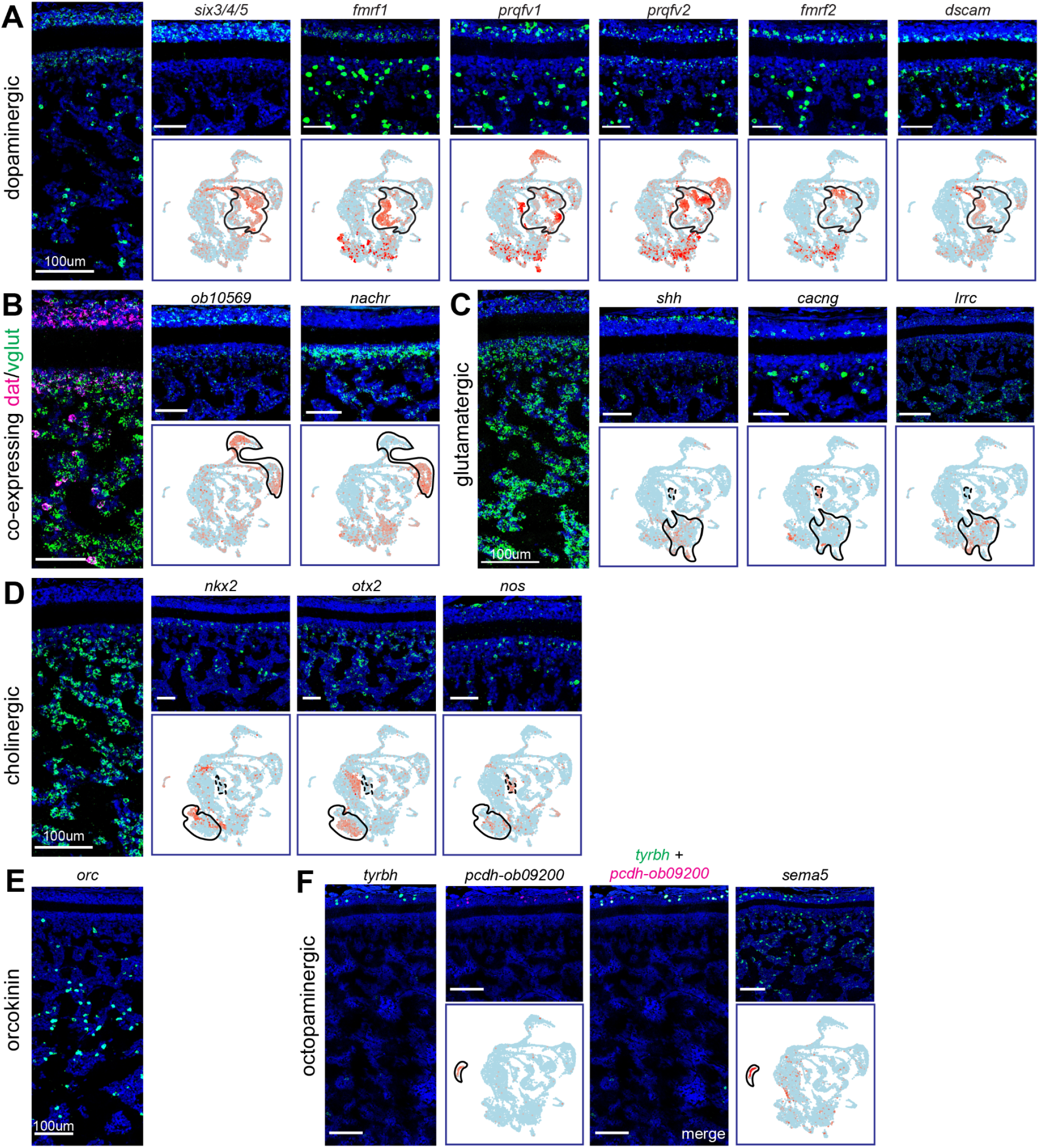
Anatomical organization of major cell classes and subtypes within the optic lobe based on scRNA-seq and FISH. (A) Dopaminergic neuron organization. *dat*+ cells are divided into two major subtypes based on the additional differential expression of either *six3/4/5*+ or *fmrf1*+. Additional markers show subtypes within dopaminergic neurons, depicted through FISH and their corresponding scRNA-seq feature plots, where the clusters that represent each of the major cell classes are circled in black. *six3/4/5*+ cells are a subset that are localized to the OGL. *fmrf1* expression is specific to the bottom sublayer of the OGL. Interspersed within *dat*+ cells are a set of four neuropeptides (*fmrf and prqfv* gene families), which are expressed in sublayers of the OGL and have stronger expression in the medulla. A subset of the *fmrf1+* population expresses *dscam* and is localized to a narrow sublayer on the border of the OGL and plexiform layer. Throughout this figure, nuclei are stained with DAPI (blue), and scale bars represent 50um. (B) Neuron organization for cells that co-express *dat* and *slc6a15/18*. Two sets of clusters have co-expression of *dat* and *slc6a15/18*. One set of these clusters is marked by an unidentified gene *obimac0010569* and is mainly localized to the OGL, whereas the other cluster is marked by expression of *nachr* and is mainly localized to the IGL. (C) Glutamatergic neuron organization. Glutamatergic neurons are found across the OGL, IGL, and medulla. The outermost sublayer of the OGL is demarcated by *shh*, the IGL contains *cacng*+ cells, and the deep medulla contains *lrrc*+ cells. (D) Cholinergic neuron organization. Cholinergic neurons are found in the IGL and medulla. While *nkx2*+ cells are in the IGL with sparse expression in the medulla, *otx2*+ cells are restricted to the medulla, and *nos*+ is mainly in the IGL. (E) Orcokinin neuron organization. *Orc*+ cells are highly specific and are found in the deeper region of the medulla. (F) Octopaminergic neuron organization. Octopaminergic neurons are identified by *tyrbh* expression and are mainly localized to a few cells in the OGL. This population also co-expresses a protocadherin. From left to right: *tyrbh, pcdh-obimac0009200*, double FISH of *tyrbh* and *pcdh-obimac0009200*, and a single FISH of *sema5*.

Expression heterogeneity of additional genes suggests these two dopaminergic cell groups can be further subdivided. The clusters within the *six3/4/5*+ group differentially express several genes encoding neuropeptides (*prqfv1, prqfv2*, and *fmrf2*), and FISH shows that these are expressed in distinct, non-overlapping sets of neurons (Fig. 4A). On the other hand, a subset of the *fmrf1+* group expresses the adhesion molecular *dscam*, which has been shown to mediate cell-type specific self-avoidance among dopaminergic amacrine cells^46^ and sublayer-specific connectivity in amacrine and bipolar cells^47^ in the vertebrate retina. It is intriguing to note that, based on FISH, the *dscam+* cells form a narrow band along the deep border of the OGL and the plexiform layer (Fig. 4A), consistent with a role in sublayer specificity. Together, these data demonstrate further heterogeneity within dopaminergic cell types in the OGL, with a spatial organization from superficial to deep layers.

#### Dopamine + glutamatergic neurons

Drawn to the co-expression of *dat* and *slc6a15/18* in both the scRNA-seq and FISH data, we next sought to further delineate this putative cell class. The scRNA-seq clusters with overlapping *dat* and *slc6a15/18* expression suggested that two prominent groups, clusters 5/6 and 37, might correspond to subtypes. Furthermore, the expression of *dat* and *slc6a15/18* significantly overlap in both the OGL and IGL, suggesting that one of the clusters might represent the former while the other defined the latter (Fig. 3A-B). The currently uncharacterized gene *obimac0010569* (see Supplemental Text for further information) and an acetylcholine receptor (*nachr*) were uniquely expressed in these two groups (Fig. 4B). FISH revealed *obimac0010569* as demarcating a broad band of expression in the OGL, thus supporting a third large population of amacrine cells within the OGL, along with the two groups of dopaminergic neurons described above. On the other hand, FISH of *nachr* corresponded to a sublayer within the IGL. It has previously been demonstrated that a set of neurons within the IGL projects axons centrifugally back to the retina^5^ and that this projection mediates retinal light adaptation in a dopamine-dependent manner^48^. It is intriguing to speculate that this circuit therefore likely corresponds to the *nachr+* dopaminergic+glutamatergic population. Moreover, *obimac0010569* is particularly striking in that it has no clear ortholog to any species outside of octopuses (see Supplemental Text for further characterization) and, therefore, may reveal octopus-specific molecular mechanisms of specialization within this cell population. Together these data support the idea that the dopamine+glutamatergic neurons consist of two distinct subtypes, one in the OGL and one in the IGL.

#### Glutamatergic neurons

We next focused on the subtypes of putative glutamatergic neurons, which includes several smaller clusters (33, 34, 28) in addition to a set of larger clusters (29-32) (Fig. 2A). Examination of gene expression in the smaller clusters revealed that these contain further subtypes. The first of these clusters (33) was defined by sonic hedgehog (*shh*), a signaling molecule that plays a role in axon guidance and patterning in the nervous system across many species^49^. FISH revealed that expression of *shh* is mainly restricted to a narrow band of neurons in the most superficial OGL, identifying yet another cellular subtype within the OGL (Fig. 4C). A second cluster (34) is marked by the voltage gated calcium channel gamma subunit 5/7 (*cacng*), and FISH demonstrated that this corresponds to a narrow band of neurons at the border of the IGL and medulla (Fig. 4C). Finally, a third cluster (28) specifically expressed a member of the leucine-rich repeat family of cell adhesion molecules (*lrrc*), which have been shown to play a role in cell-type specific synaptic connectivity in fly and mammal nervous systems^50^. FISH showed that this group was localized to the deeper region of the medulla (Fig. 4C). Upon investigating markers expressed in the larger clusters (29-32), we found that a subset of neurons express *vat1*, which FISH demonstrated to also be localized to cell bodies in the deeper region of the medulla (see Fig. 7F). Thus, the glutamatergic neurons primarily constitute a large population of cells within the medulla along with two additional subtypes with highly specific sublayer localization with the outer OGL and inner IGL respectively.

#### Cholinergic neurons

The last large neurotransmitter population is the putative cholinergic neurons. scRNA-seq data revealed a large population of cells (clusters 7-9) and a much smaller population (cluster 35) that both express *vacht*, a marker for cholinergic transmission. FISH for *vacht* shows that cholinergic neurons are located throughout the medulla, with more restricted expression in IGL (Fig. 3B, 4D). Moreover, the scRNA-seq data revealed that clusters that comprise the larger population can be delineated based on the expression of family-members of two homeobox transcription factors, *nkx2* and *otx2*. FISH data for these two markers demonstrated that the *nkx2+* population is located in the IGL and superficial radial columnar region of the medulla, while the *otx2+* population is not expressed in the IGL and, instead, is found throughout the medulla (Fig. 4D). In exploring the scRNA-seq data, we found that the *nkx2+* cluster also expresses a protocadherin (*obimac0026462*), and FISH confirms that *obimac0026462+* cells are expressed both in the IGL and medulla (see Fig. 7E).

Intriguingly, we observed that neurons in the smaller cluster (35) selectively express nitric oxide synthase (*nos*), suggesting that they represent a distinct cell type that uses nitric oxide as a signaling molecule in addition to acetylcholine. Correspondingly, FISH revealed that these neurons form a narrow sub-layer within the superficial IGL (Fig. 4D). Interestingly, scRNA-seq data show that *nos+* cells and the glutamatergic *cacng+* cells, both in the IGL, also express an unidentified gene *obimac0022194* (see Supplemental Text). A FISH of *nos, cacng*, and *obimac0022194* together demonstrate that these three genes are expressed most prominently in the IGL and both *nos* and *cacng* are co-expressed with *obimac0022194* (data not shown), suggesting these clusters may have some shared function based on this unidentified gene..

Thus, the cholinergic neurons constitute a large population of IGL/medulla cells with distinct anatomical positions defined by a small sub-layer of *nkx2+* cells in the IGL and superficial medulla, *otx2+* cells throughout the medulla, and *nos+* cells mainly in the superficial IGL.

#### Orcokinin and Octopaminergic neurons

Next, we sought to determine the identity of the two remaining putative neuronal clusters (37 and 4), which were defined by highly specific expression of *orc* and *tyrbh*, respectively (Fig. 2A-C). Cluster 37 does not express any of the neurotransmitter markers we assessed, but is demarcated by the expression of the neuropeptide *orc*, and FISH revealed that *orc*+ cells are a sparse, scattered population throughout the deeper region of the medulla (Fig. 4E). The final cluster of mature neurons, cluster 4, selectively expressed a number of genes, including *tyrbh*, the synthetic enzyme for octopamine, generally considered to be the invertebrate analog of norepinephrine^51^, both of which play a role in arousal and other aspects of behavioral state across species. FISH identified a discrete population of octopaminergic *tyrbh*+ neurons in the OGL (Fig. 4F). Among the other genes that are unique to this cluster is a protocadherin family member (*obimac0009200*), which co-expresses with *tyrbh* restrictively in the OGL (Fig. 4F), and semaphorin-5 (*sema5)*, which FISH validates as being highly expressed in the same region of the OGL along with some expression throughout the medulla (Fig. 4F). Thus, this cluster also expresses genes that serve as adhesion molecules (protocadherin *obimac0009200*) and axon guidance cues (*sema5*) in the visual system of other species^52,53^. As a counterpart to these octopaminergic neurons, scRNA-seq data reveal that the putative octopamine receptor (*obimac0000086, octr*) is found in the cholinergic and glutamatergic medulla clusters (data not shown), suggesting that their downstream synaptic targets may be cells in the medulla.

### Immature Neurons

Next, we examined the putative immature neuronal clusters that were identified in the scRNA-seq data based on lower expression of *syt* and the absence of mature neurotransmitter markers. These clusters make up 31% of the putative neurons in our data. We found that the clusters of these cells were organized with one main large cluster and several unique cluster ‘arms’ that may represent discrete populations of developing neurons associated with mature cell types (Fig. 2A). Intriguingly, one unidentified gene (*obimac0011980*) appears to comprehensively encompass all immature clusters (Fig. 5A-B, see Supplemental Text for further characterization). The immature clusters can then be segregated into three subgroups that are demarcated by complementary expression of genes tumor necrosis factor receptor (*tnfr;* clusters 18-20, 27), myoneurin (*mynn;* clusters 22-24), and big brain (*bib;* clusters 25-26) (Fig. 5A-B), all of which are known to play a role in neural development in other species. Since these three genes segregate the population of immature neurons, we used FISH to confirm their expression *in vivo* and examine their relationships to the location of mature cell types described above (Fig. 5C).

**Figure 5.**
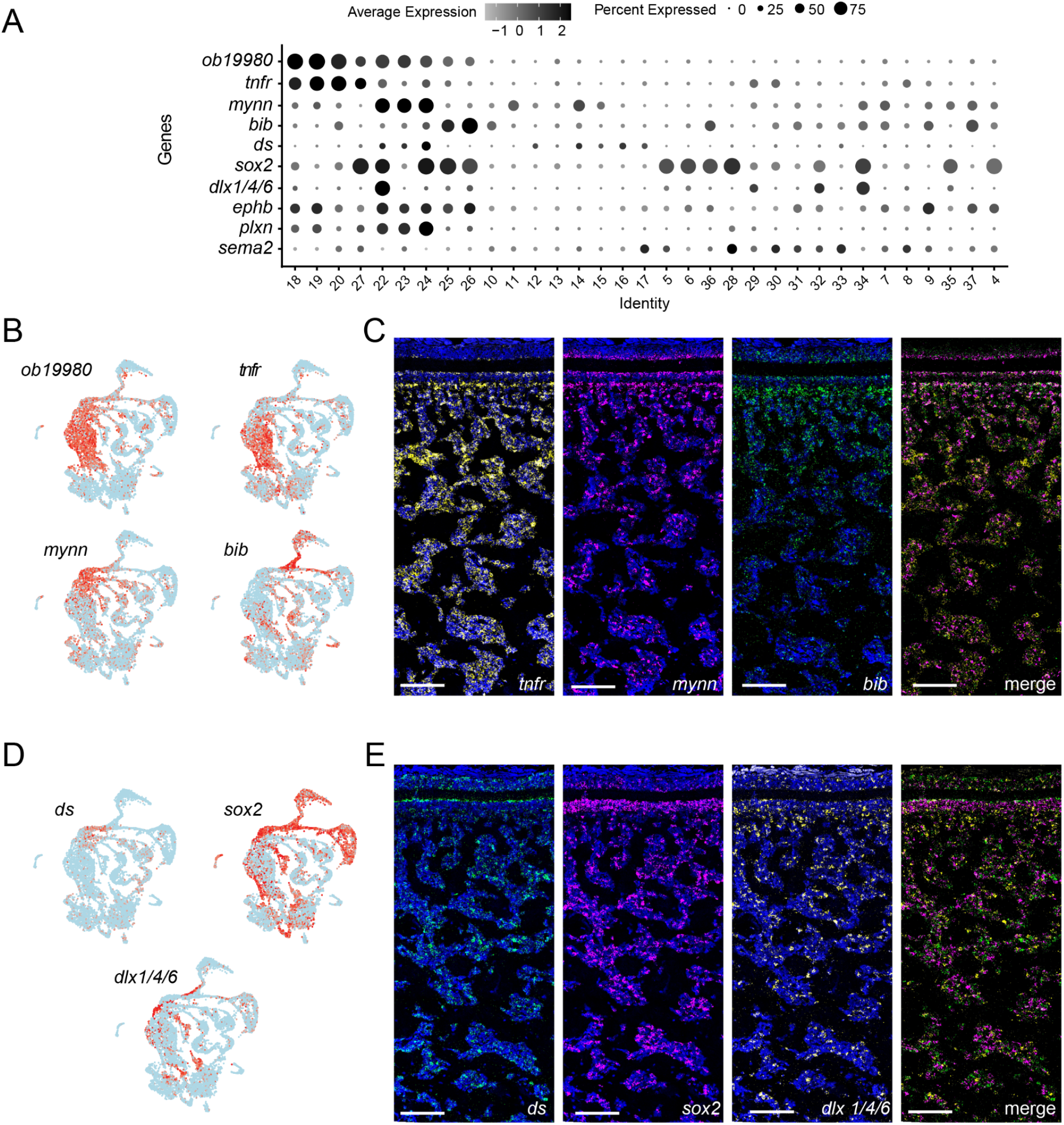
Gene expression and spatial organization of putative immature neurons. (A) Dot plot of genes expressed in immature neurons. (B) Feature plot of uncharacterized cephalopod-specific gene *obimac0019980*, which demarcates the putative immature neuron clusters, as well as three genes that define distinct subgroups within the immature neurons: *tnfr, mynn*, and *bib*. (C) FISH of the genes delineating the three subgroups shown in B. From left to right: *tnfr* is found mostly in the medulla; *mynn* is most prominent bordering the PL; *bib* is expressed mostly in OGL, IGL, and the most superficial layer of the medulla; merged FISH. Throughout this figure, DAPI is shown in blue on individual FISHs, and scale bars indicate 100um. (D) Feature plots of additional markers from development-related gene family trees demonstrating further cell type diversity. (E) FISH showing anatomical organization of the genes shown in D. From left to right: *ds, sox2, dlx1/4/6*, and a merged FISH of the three genes.

The receptor *tnfr* has been extensively implicated in brain development due to its role in regulating neuronal growth and differentiation^54^. *tnfr* expression appears in scRNA-seq clusters that are transcriptionally similar to mature clusters 8 and 30, which are presumed to represent the medulla. Interestingly, FISH data for the sub-group marker *tnfr+* revealed expression in cells that are more abundant throughout the medulla compared to the other putative immature subtypes. On the other hand, scRNA-seq data shows that *mynn*, a zinc finger protein family member associated with neuromuscular synapse formation in mice^55^, is expressed in a number of clusters in the “arms’’ leading to mature cell types for dopaminergic neurons of the OGL. Correspondingly, FISH data shows that the *mynn+* cell types border the plexiform layer with some cells in the medulla. Lastly, *bib*, a known neurogenic molecule in *Drosophila*^*56*^, expresses in “arms” leading to the clusters that correspond to the two prominent clusters of dopaminergic+glutamatergic neurons, and *bib* was expressed most strongly, though not exclusively, in cells along the bottom borders of the IGL and OGL. Notably, although these three groups each show highest expression in the regions corresponding to mature types, they are also found throughout the medulla, which may represent the ongoing migration of immature neurons into the optic lobe ^57^. Altogether, these three immature cell types form complementary patterns of expression from the deeper central tangential layers of the medulla (*tnfr+*) to more superficial layers of the optic lobe (*mynn+* and *bib+*), corresponding to locations of distinct mature clusters, and supporting the hypothesis that these represent three populations of immature neurons that may give rise to corresponding mature populations of the optic lobe.

Furthermore, a number of additional known developmental genes (Fig. 5A, 5D) were expressed in the immature clusters in the scRNA-seq data, and we confirmed three of these genes using FISH (Fig. 5E). Dachsous (*ds*), a cadherin that plays a role in non-canonical Wnt signaling in other species^58^, was expressed within the *bib+* clusters, and FISH confirmed that its expression similarly localized mostly to the borders of OGL and IGL. *sox2*, a homeobox gene involved in cell fate in both vertebrate retina and visual thalamus^59,60^, had scRNA-seq expression across multiple immature and corresponding mature clusters, including the two dopaminergic+glutamatergic classes and neurons of the medulla. FISH data showed corresponding expression in the OGL and IGL, as well as sparsely throughout the medulla. Finally *dlx1/4/6*, a homeobox gene, was expressed in several clusters within the immature neurons. FISH showed an intriguing expression pattern in the optic lobe, with strongest expression in the IGL and superficial radial columnar region of the medulla, consistent with a superficial to deep patterning reminiscent of the role of *dlx1/4/6* in proximal-distal patterning in other species^61^.

In addition to the cluster-specific markers described above, we identified three genes that have well established roles in patterning the nervous system in vertebrates and other invertebrates^52,62^ - *ephb, sema2*, and *plxn* - which had complementary expression patterns to each other across the scRNA-seq data (Fig. 6A). Graded expression of Eph receptors and their ephrin ligands, and plexins and their semaphorin ligands, play important roles in establishing large-scale organization during development, including topographic map formation^52,62^. Notably, these genes had distinctive patterns of expression across the optic lobe (Fig. 5A, 6B). *ephb+* showed a gradient of expression, restricted to the IGL and the superficial radial columnar region of the medulla. *sema2+* had an opposite gradient, with strongest expression in the deeper central tangential region of the medulla, while *plxn+* had distinctive expression along the borders of the OGL and IGL along with sparse expression throughout the medulla. These graded expression patterns suggest that these genes may play similar roles in large scale patterning of the octopus visual system as well.

**Figure 6.**
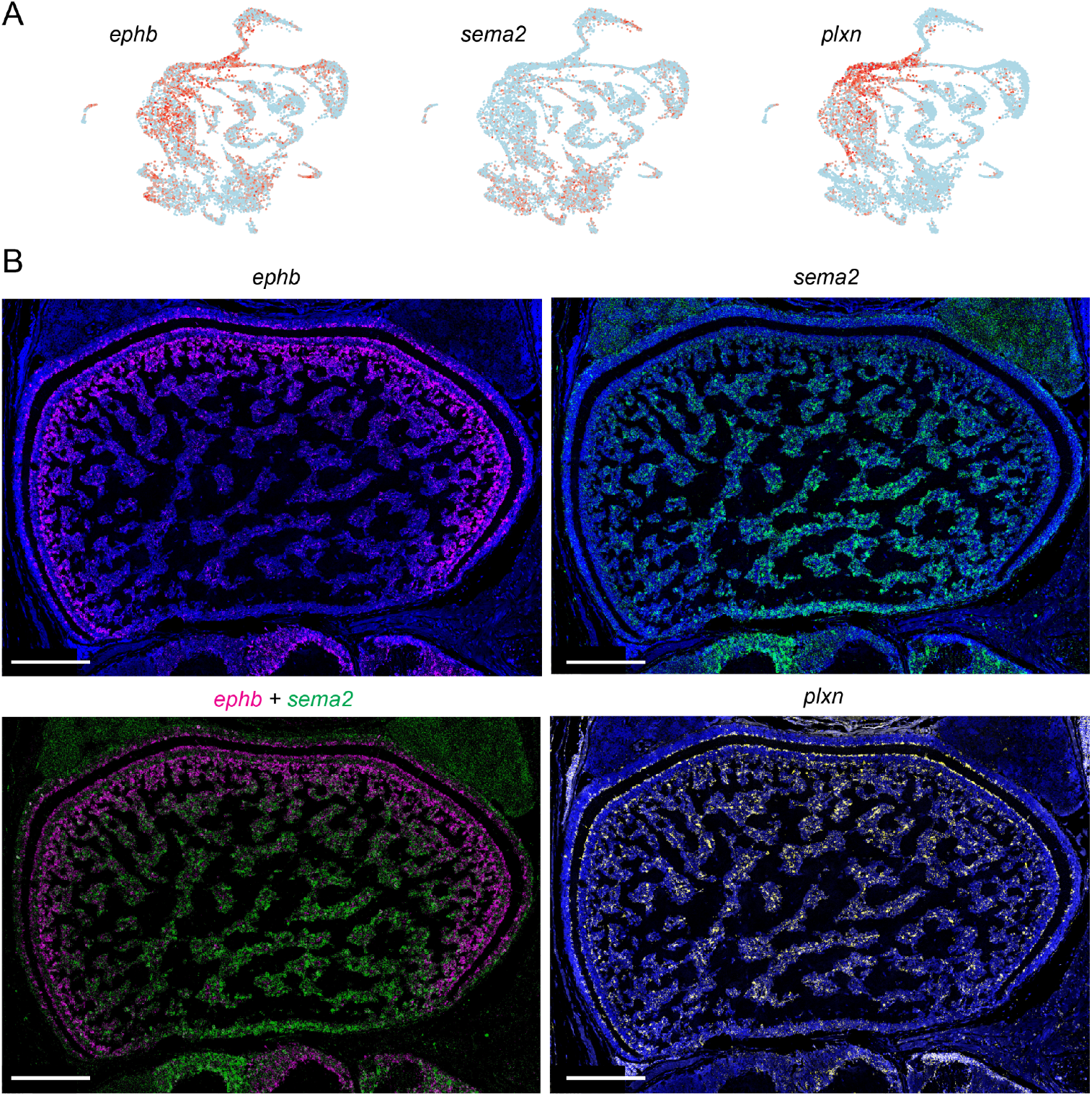
Established patterning molecules are expressed in the optic lobe. (A) Feature plots showing scRNA-seq support for genes that are well-known for patterning in other organisms. (B) FISH showing gradients of expression for the three genes depicted in A, with the bottom left image showing a merged image of *ephb* and *sema2*. Nuclei are stained with DAPI in blue. Scale bars indicate 200 um.

Together, these data demonstrate that these putative immature neuronal classes are found in distinct anatomical locations with discrete subtypes that can be molecularly defined. Moreover, conserved families of genes, as well as novel cephalopod- and octopus-specific genes, define unique categories, tantalizingly suggesting the possibility of both evolutionarily conserved and lineage-specific molecular mechanisms for development and function.

### Cell-type and sub-layer organization of mature neurons in the optic lobe

A driving goal of this project was to identify the “parts list” of the optic lobe and create an integrated model of cell type organization within the octopus visual system. We therefore incorporated the findings for the mature neurons into a schematic spatial map of the optic lobe (Fig. 7A-B). This represents a comprehensive molecular description of neural subtype organization of the optic lobe to complement the anatomical description provided by Young^5^, and reveals detailed cell type-specific structure in each of the major regions of the optic lobe. Here we summarize this organization and present multiplexed FISH data for markers within each layer to explicitly demonstrate the sub-layer organization.

**Figure 7.**
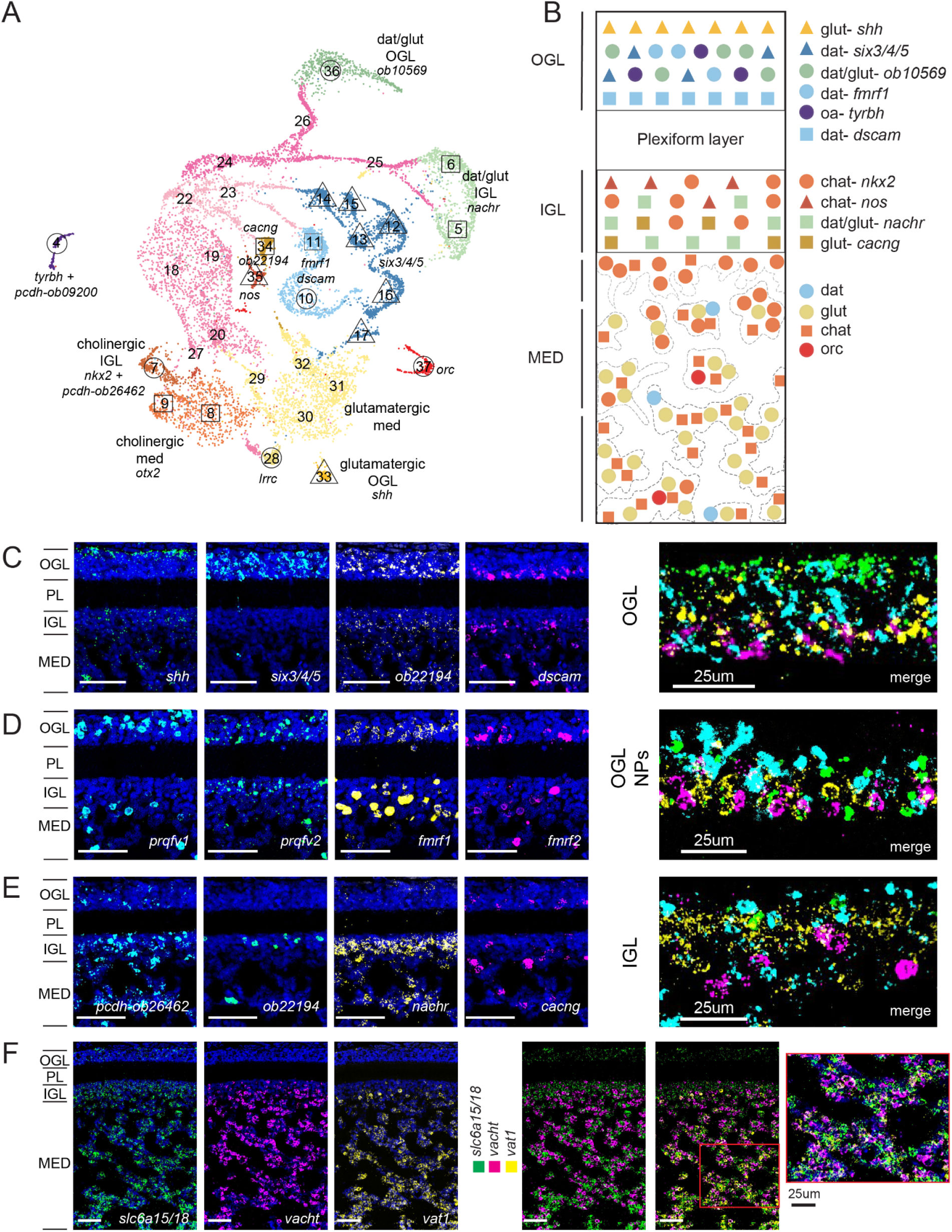
Summary of mature neuronal circuitry of the optic lobe. (A) UMAP showing cell subtypes in each neuronal class, along with annotation of spatial localization within the optic lobe. (B) Schematic of cell type organization of the optic lobe. (C) FISH showing sublayers of the OGL. Sublayers of OGL are demarcated from most superficial to deepest in order by expression of *shh, six3/4/5*, unidentified gene *obimac0010569*, and *dscam*. A merge of these is shown in the last panel. Throughout this figure, nuclei are stained with DAPI in blue, and, unless otherwise noted, scale bars represent 50um. (D) FISH of neuropeptides that subdivide dopaminergic neurons in the OGL. In order from most superficial to the deepest: *prqfv1, prqfv2, fmrf1*, and *fmrf2*. (E) FISH showing sublayers of IGL based on expression of *pcdh-obimac0026462* (specific to *nkx2+* cluster), unidentified gene *obimac0022194* (specific to *nos+* cluster), *nachr*, and *cacng*. (F) FISH showing organization of the medulla. The medulla has cell body islands with non-overlapping expression of populations of glutamatergic (*slc6a15/18+*) and cholinergic (*vacht+*) neurons, while *vat1* is expressed in a subset of these populations. A double FISH of *slc6a15/18* and *vacht* shows segregated cell body islands of glutamatergic and cholinergic neurons in the medulla, and a triple FISH including *vat1* shows that expression of these genes spans a majority of medulla cell bodies. The red box denotes the inset shown in the previous panel.

#### Outer granular layer

We find four broad groups of cell types within the OGL (Fig. 7C). First, a cluster of glutamatergic cells lines the most superficial aspect of the OGL, which also expresses *shh*. Second, a cluster of dopaminergic+glutamatergic neurons is located in the central OGL, marked by an uncharacterized gene *obimac0010569*. Third, we identify a specific subset of octopaminergic neurons that co-express *tyrbh* and *pcdh-ob09200*. Finally, a diverse group of dopaminergic-only neurons span sublayers of the OGL. This group falls into two major divisions: *six3/4/5*+ in the central OGL and *dscam+* in the deep OGL. Notably, the expression of two pairs of *prqfv* and *fmrf* neuropeptides (NPs) within the *dat+* group also shows a progression across the depth of the OGL (Fig. 7D). Based on Young’s finding that neurons of the OGL have an amacrine morphology, this great diversity is reminiscent of the large diversity of amacrine cells in the vertebrate retina ^63^.

#### Inner granular layer

We also find significant diversity within the IGL (Fig. 7E). The largest population of cells therein consists of dopaminergic+glutamatergic neurons, which form a distinct band as identified by expression of *nachr*. In addition, a small population of cholinergic neurons expressing *nos* lines the superficial IGL (*obimac0022194*+ is shown in Fig. 7E as proxy for *nos+* cells), while a sub-group of glutamatergic neurons (*cacng*) lines the deep IGL. In addition, the *nkx2+* cholinergic group spans from the IGL into the medulla (Fig. 7E includes *obimac0026462* in place of *nkx2*). We find that the IGL consists of at least three neuronal cell types with distinct sub-layer expression patterns.

#### Medulla

The medulla comprises a majority of the optic lobe and largely consists of two distinct populations of glutamatergic and cholinergic neurons (Fig. 7F), with glutamatergic neurons more prevalent in the deeper central tangential region of the medulla. Notably, these are intermingled within the cell body “islands” of the medulla, suggesting that, at least at the level of these two large populations, there is no functional segregation across the islands. However, there are apparent distinctions in gene expression across the depth of the medulla. Within the cholinergic group, *nkx2+* neurons are more superficial in the radial columnar region, while *otx2+* neurons are located throughout the medulla (Fig. 7B). In addition, there is a gradient of *vat1* expression that is shared across the cholinergic and glutamatergic neurons, with the strongest expression of *vat1+* cells found in the deeper central tangential region of the medulla (Fig. 7F). Finally, there are sparse populations of neurons within the central tangential region of the medulla that express extremely high levels of neuropeptides, including the two sets of *fmrf* and *prqfv* related genes and *orc* (Fig. 4A, E). It is possible that these represent neurons projecting to downstream brain regions where these neuropeptides may play a role in regulating behavioral outputs^14,18,20,21,64^.

Together, this description of mature neuron identities and their spatial organization shows significant diversity of cell types within each of the neurotransmitter classes and demonstrates an intricate sub-layer organization across major subdivisions of the optic lobe. This parts list and corresponding spatial layout of the mature circuitry provide a basis for studying the visual processing performed by the optic lobe.

## Discussion

By combining scRNA-seq data mapped onto an updated genome with RNA FISH, we both identified transcriptional cell types of the octopus visual system and determined the spatial organization of cells with unique molecular signatures within the optic lobe. In this single-cell atlas we reveal six major cell classes of mature neurons based on neurotransmitter types in addition to groups of non-neuronal and developing cells. We demonstrate that additional genes further identify subtypes within the neurotransmitter cell classes, and these genes include transcription factors, adhesion molecules, developmental signaling molecules, and genes with unknown identity that may be unique to cephalopods. Furthermore, these transcriptional cell types are located within discrete locations of the optic lobe, identifying previously unknown cell-type diversity and sub-layer organization. This study is the first to comprehensively delineate the cell types of the octopus optic lobe at the molecular level. It thereby contributes to a recently growing literature on transcriptomics of the cephalopod nervous system^65–67^, and lays the basis for the investigation of the role of distinct cell types in visual processing, as well as the development of tools to target specific cells based on their molecular signatures.

In addition to revealing the overall molecular architecture of the optic lobe, a number of the specific findings shown here have implications for function and development of the octopus visual system, particularly from a comparative perspective. We find a wide array of cell types within the OGL, which was previously shown to consist of amacrine cells. Among the three broad classes of amacrine cells in the OGL that Young described^5^, we find at least eight clusters of specific cell types that all have distinct spatial localization within sublayers of the OGL (Fig. 7C, D). These cell types are defined by neurotransmitters (largely dopaminergic, but also exclusively or co-expressing glutamate), *fmrf* and *prqfv* neuropeptides, a transcription factor (*six3/4/5*), an adhesion molecule (*dscam*), and a developmental signaling molecular (*shh*). This array of cell types bears a strong resemblance to the diversity of amacrine cells in the vertebrate retina, as demonstrated by a recent single-cell gene expression study in mice^63^, where over 60 amacrine cell types were identified based on expression of different neurotransmitters and neuropeptides. However, in contrast to the vertebrates, where amacrine cells primarily express the inhibitory neurotransmitters GABA or glycine^68^, in the octopus we find that OGL neurons are predominantly dopaminergic. Notably, though, there is a specific population of dopaminergic amacrine cells in the vertebrate retina^69^, and it has been shown that dendritic tiling in these amacrine cells is dependent on *dscam*^*46*^, which strikingly is also expressed in a subset of dopaminergic OGL cells here. Moreover, in the vertebrate retina, distinct amacrine cell classes have been linked to a range of specific visual computations, including luminance adaptation, direction selectivity, and local versus global motion processing^70^. It will be intriguing to see whether similar functions can be assigned to the diversity of cells we find within the OGL based on the markers we have identified.

We also identify a sparse but highly distinct population of neurons in the OGL that express *tyrbh*, the synthetic enzyme for octopamine, and the corresponding receptor *octr* expressed throughout the larger scRNA-seq clusters that reflect cells in the medulla. In both flies and mice, locomotion and arousal have profound effects on visual processing^71–73^, which, in flies, are mediated by octopamine^74^, and in mice are mediated in part by norepinephrine^75^, considered the vertebrate analog of octopamine^51^. Strikingly similar impacts of arousal on visual responses in octopuses were observed in an early study in the optic lobe, as measured by EEG^76^, suggesting a potential role for this octopaminergic system. Directly linking cell types with such hypothesized roles in visual processing is an exciting future prospect stemming from these findings.

Another important finding presented here is the delineation of a large population of immature neurons, and their corresponding gene expression patterns, within the octopus visual system. *O. bimaculoides* are born capable of feeding and living independently from hatching, relying on a variety of visually guided behaviors^29,30^. Despite their precociousness at birth, octopuses continue to grow exponentially in size throughout their life^77^, and this includes growth within the optic lobes and other brain regions^78,79^. At the stage we study them, 1.5 months, they are ∼10x times larger in mass than at hatching, but will continue to grow up to 500x that size by 1 year of age, which is their typical lifespan^77^. Hence, the finding of such a large population of immature neurons is not surprising given the octopus’s need to generate additional neurons to support such massive brain growth. The fact that these immature neurons need to integrate into a functioning visual system raises important questions about how neuronal morphogenesis, synapse formation, and refinement are coordinated to allow them to integrate without disrupting ongoing function. The visual systems of fish and birds also continue to grow throughout their lifetime, but coordinated growth of the retina and downstream brain areas occurs by expansion along the edge in a proliferative marginal zone^80^. By contrast, interestingly, we find that the immature neuronal population in the octopus is distributed across the optic lobe. It is currently unknown how the cephalopod retina itself grows through adulthood, whether through spatially restricted marginal zones or through continued addition of new photoreceptors across the entire retina. Future work examining the ongoing developmental expansion of the optic lobe may reveal how new neurons coordinate their integration into the fully functioning visual system of the growing octopus.

We also identify subtypes within the immature population that are associated with mature cell types, suggesting different developmental trajectories and progenitor populations. We find genes associated with both the immature population and mature cell types that may contribute to their identity and eventual function. Some are members of gene families with well known developmental function, such as *shh* known in vertebrates for patterning neural progenitors, while others are genes unique to cephalopod or octopus lineages. The expression of genes that delineate sub-layers within the optic lobe is suggestive of both conserved and unique gene functions in building the octopus visual system. For example, *shh* defines a narrow layer of cell bodies in the most superficial OGL, whereas *dscam* demarcates the deeper OGL. Additionally, a specific protocadherin, *obimac0009200*, marks the octopaminergic OGL population, suggesting a potential role in establishing its connectivity. We also find genes that delineate broad population of immature neurons, including a cephalopod-specific gene of unknown identity (*obimac0011980)* that is expressed across all immature neuron clusters, and the neurogenic signaling molecule *bib*, that is associated with immature neuron clusters leading to the two mature populations expression *dat* and *vglut*, suggesting a role in specifying the dopaminergic+glutamatergic population. Another striking developmental finding is the presence of complementary gradients in *ephb* receptor and *sema2* expression in the optic lobe, suggesting that these may play a similar role in setting up the spatial organization of the visual system as they do in flies and vertebrates. Causal tests of the role of all these molecules will likely depend on the continued development of technology for knocking down or manipulating gene expression in cephalopods, as in the recent use of CRISPR to knock out pigmentation in squids^81^, or the use of pharmacological approaches to target known signaling pathways, as was recently used to test the role of Wnt signaling in cephalopod lens development^82^.

### Implications for future studies

We note that although we have delineated a number of subtypes within the major neurotransmitter populations of cell types in the octopus optic lobe, there is almost certainly additional diversity to be explored in future studies. For instance, the large clusters of the medulla could be further subdivided based on analyses focusing specifically on this population. In fact, even some of the existing cluster divisions we identified here have not been uniquely identified via FISH due to the lack of specific markers. However, our goal here was to provide the broad organizational structure of the octopus visual system as a conceptual framework for future studies. Indeed, studies of the vertebrate retina have progressed from the basic delineation of five major cell types (photoreceptors, horizontal cells, bipolar cells, amacrine cells, and retinal ganglion cells) to our current understanding of the tremendous diversity within each of these types, including 30+ cell types within the retinal ganglion cells alone^83^. The initial description of cell types for the octopus visual system that we provide here can serve as a basis for delving further into such diversity.

In this study we defined cell types based on gene expression and related these cell types to their anatomical location, revealing the molecular architecture of cell types within the octopus optic lobe. Future studies that characterize these cell types by relating them to other key aspects of neural identity, including anatomical morphology based on single cell labeling, downstream projection targets based on retrograde tracing, and visual response properties based on calcium imaging^84^, will be crucial in decoding cephalopod visual function. The atlas we present here provides a roadmap for such studies, and more generally provides a path forward towards cracking the functional, developmental, and evolutionary logic of the cephalopod visual system.

## Acknowledgements

We thank members of the Niell, Miller, and Kern labs for helpful discussions and comments on the manuscript. We acknowledge support from the University of Oregon Genomics & Cell Characterization Core Facility and Imaging Core Facility. We thank Leah Deblander for assistance with slide-scanner imaging and Poh-Keng Loi for performing tissue sectioning. This work was supported by National Institutes of Health R01NS118466-01 (C.M.N.) and R01NS118466-01S1 (C.M.N., J.O.SC.), Office of Naval Research N00014-21-1-2426 (C.M.N.), University of Oregon Renee James Seed Grant (A.C.M., C.M.N), National Science Foundation Graduate Research Fellowship Grant No. 1842486 (G.C.C.), and R01HG010774 (A.D.K).

## Author Contributions

C.M.N. and A.C.M. conceived and oversaw the project. J.O.SC., D.M.P., and J.R.P. collected sequencing data. G.C.C. and A.D.K. performed genome assembly and annotation, J.R.P. performed cross-species gene identification, J.O.SC., A.C.M., and C.M.N. performed scRNA-seq analysis. J.O.SC. and D.M.P. performed FISH experiments. All authors contributed to writing and editing of the manuscript.

## Methods

### Animal use

*Octopus bimaculoides* were obtained from the Cephalopod Resource Center at the Marine Biology Laboratory (Woods Hole, MA) and from Aquatic Research Consultants (San Pedro, CA). They were kept at the University of Oregon in a closed circulating 250 gallon aquarium system in artificial seawater, and fed daily on a rotating diet of frozen shrimp, crabs, and fish. All husbandry and experimental protocols were in accordance with the EU 2010/63/EU^85^ and AALAC guidelines for the use and care of cephalopods for research.

### Genome sample collection and sequencing

Optic lobe tissue was dissected from an adult female *O. bimaculoides* for whole genome sequencing. Tissue was sent to the University of Oregon Genomics & Cell Characterization Core Facility (GC3F) for DNA extraction and sequencing. High molecular weight genomic DNA was extracted using a Nanobind Tissue Big DNA kit (Circulomics). A Pacific Biosciences standard HiFi library was prepped with a SMRTbell Express Template Prep Kit 2.0. Genomic DNA was sheared at 20kb target size with a Megaruptor 2 instrument (Diagenode). BluePippin size selection (Sage Science) was used to omit the smallest fragments (<10-14kb) to enrich for longer fragments. Two HiFi genomic circular consensus sequencing (CCS) SMRTbell libraries were prepared as input for five HiFi SMRT cells. Single molecule sequencing of both libraries was conducted with a PacBio Sequel II system. After sequencing, data was imported into SMRT Link to generate 5.8 million HiFi reads with the CCS algorithm and create fastq files.

Four tissues were dissected from 6 week old juvenile octopuses to be used in Iso-Seq sequencing: optic lobe, central brain, retina, and arm. RNA extractions were performed using a RNeasy Plus Mini Kit (QIAGEN). A single bulk non-barcoded SMRTbell Iso-Seq library was prepared according to the manufacturer’s protocol (PacBio) by GC3F. A multiplexed Iso-Seq library was sequenced across a single PacBio Sequel II SMRT cell. IsoSeq3 in SMRT Link was used to generate fastq files containing 1.08 million full-length transcripts.

### Genome re-assembly and annotation

We used HiFiASM v0.15.5-r352^86^ to assemble a contig-level genome with HiFi reads as input and default parameters. After initial assembly, duplications were removed using Purge_dups ^87^. Protein-coding genes were annotated using existing bulk RNA sequences and our newly generated Iso-Seq data. Bulk RNA data was aligned to the genome assembly using Hisat v2.2.1 ^88^ and gene predictions were assembled with StringTie v2.1.6 ^89^ using parameters `-c 4 -m 200 -j 3`. All other parameters were set to default. To generate gene predictions with Iso-Seq data, we mapped the full length, non-chimeric reads (FLNC) to the genome using minimap2 (cite) using parameters `-ax splice -uf --secondary=no -C5 -O6,24 -B4.` After alignment, cDNA_cupcake (https://github.com/Magdoll/cDNA_Cupcake) was used to collapse alignments into transcript models. Unique transcripts with degraded 5’ ends were filtered out of the final annotation file with `filter_away_subset.py`. StringTie and cDNA_cupcake annotations were combined using TAMA merge ^90^ with parameters `-e longest_ends -d merge_dup`.

The transcripts of the resulting gtf were used to run blastp against a Uniprot database and to run hmmer against the pfam database. The resulting hits were used as input for Transdecoder v5.5.0 (https://github.com/TransDecoder/TransDecoder) to predict single best coding regions. The existing mitochondrial genome and annotation were concatenated to the assembled genome and annotation files, respectively. This resulted in a final number of 18,896 gene annotations.

Orthologous relationships between our predicted genes and those of distant species were identified using Orthofinder v2.5.2^34^. We used default parameters to cluster sequences into orthologue groups using sequences from eight species including *Homo sapiens* (hg38), *Mus musculus* (GCA_000001635.9), *Drosophila melanogaster* (GCA_000001215.4), *Aplysia californica* (GCA_000002075.2), *Crassostrea gigas* (GCA_902806645.1), *Octopus sinensis* (GCA_006345805.1), *and Sepia pharaonis* (GCA_903632075.3). Genes that did not have predicted orthogroups were also manually annotated using NCBI BLAST to assign putative identity based on homology to deposited sequences in other species.

### Cell dissociation for scRNA-seq

Animals used for scRNA sequencing were 6 week old juveniles with mantle lengths of 6.5mm-9.0mm. *O. bimaculoides* optic lobes were dissected on ice in Leibovitz-15 medium (Gibco) supplemented with 400mM NaCl, 10mM KCl, 15mM Hepes, 200 U/mL penicillin, and 0.2 mg/mL streptomycin). Single cell dissociation was performed by incubating tissue in papain (1 mg/ml; Worthington Biochemical Co) plus 1% DNaseI (10mg/ml in HBSS) in supplemented L-15 medium for 10 min at RT. The cells/tissue were gently pipetted up and down several times to dissociate large chunks. The cells/tissue were incubated for another 10 min at RT, pipetted up and down several times, and quenched in wash solution containing 2.5M glucose, 5mM Hepes, and 5% FBS in CMFSS (12mM Hepes, 435mM NaCl, 10.7mM KCl, 21mM Na2HPO4, 16.6mM glucose). Dissociated cells were passed through a 40 μM cell strainer (Fisherbrand), washed again, and resuspended in L-15 medium. A final sample cell concentration of 2000 cells per microliter, as determined on a BioRad TC20 cell counter, was used for cDNA library preparation.

### Single-cell cDNA library preparation

Sample preparation for two biological replicates was performed by the University of Oregon Genomics and Cell Characterization core facility (https://gc3f.uoregon.edu/). Dissociated cells were run on a 10X Chromium platform using 10x v.3 chemistry targeting 10,000 cells. Dissociated samples were prepared in tandem, on the same day. The resulting cDNA libraries were amplified with 11 cycles of PCR and sequenced on either an Illumina Hi-seq or an Illumina Next-seq.

### Computational analysis

The resulting sequencing data were analyzed using the 10X Cellranger pipeline, version 3.1.0 (Zheng et al., 2017) and the Seurat (version 3.1.4; Satija et al., 2015) software package for R, version 4.1.2, using standard quality control, normalization, and analysis steps.

Briefly, raw data from each biological replicate were read into R, with a minimum threshold of 3 cells and 500 genes. After visualizing the raw reads, genes, and mitochondrial percentage, we set thresholds of counts between 1000 and 20000, above 600 genes, and less than 6 percent of mitochondrial content for downstream processing.

For normalization and integration, we followed the guidelines provided by ^91^ and ^33^ respectively. We selected integration features within each dataset and applied SCTransform normalization before integrating the datasets based on Canonical Correlation Analysis^33^. For generating cell type clusters, we ran all of the following analyses on the “integrated” assay, but performed differential expression analysis on the “SCT” assay. Following standard downstream processing steps, we ran principal component analysis and UMAP on 25 dimensions. We ran FindNeighbors (dims 1:25) and FindClusters (resolution 0.85). We then generated a dendrogram and renumbered clusters based on this output. We identified the top differentially expressed markers and used these data to identify and subset the putative neurons. We also excluded one cluster from the rest of the analyses due to low number of transcripts and genes, suggesting this cluster did not represent real cellular expression. We re-ran UMAP on the subset of neurons and used this output for visualization and further cell type identification based on top differentially expressed markers. All UMAPs, including feature plots, are shown with datasets that are downsampled to 500 cells for visualization purposes. However, all dot plots show gene expression for full datasets. Feature plots shown in the figures include modified expression for visualization purposes (see code for modified and full expression). Feature plots for Fig. 2B are shown with a min.cutoff of 0 and a max.cutoff of 1, except for *syt* which has a min.cutoff of 1 and a max.cutoff of 2. Feature plots for Fig. 4 are shown with a min.cutoff of 0 and a max.cutoff of 4. Feature plots for Figs. 5-6 are shown with a min.cutoff of 0 and a max.cutoff of 2.

### RNA Fluorescence In Situ Hybridization

Tissue collection for RNA fluorescence in situ hybridization consisted of juvenile octopuses (∼4-6 weeks in age, 6-8mm in mantle length), which were first anesthetized in 4% EtOH in Artificial Seawater prior to fixation. Anesthetic replaced the seawater in the octopus’ home chamber, and the chamber was placed on ice until the octopus was no longer ventilating or responsive. Both optic lobes were dissected from each animal and immediately placed into 10% Neutral Buffered Formalin. The optic lobes were fixed for 24 hours at room temperature before being processed and embedded in paraffin and sectioned into 7um slices.

Custom probes were designed and ordered through Advanced Cell Diagnostics (ACDBio) (Hayward, CA). We followed the protocol available for ACDBio RNAscope^92^, with minor changes to optimize it for use in paraffin-embedded octopus tissue. Briefly, we first removed the paraffin through baking, xylenes, and ethanol washes. We then fixed the tissue for 30 minutes in formalin at room temperature before dehydrating the tissue with an ethanol series. We proceeded with hybridization and target retrieval: 10min pre-treatment of H2O2, 12min target retrieval in a pressure cooker, and protease plus for 25min at 40C. Slides incubated with probes for 2 hours before going into washes and 5X SSC overnight. On Day 2, we proceeded with amplification and used the appropriate HRPs and opal dyes before adding the HRP block. For multiple probes, additional HRP conjugates were added in a series-wise manner (HRP, opal dyes, block) before slides were mounted with DAPI and ProLong Gold Antifade. Slides were imaged with a Leica SlideScanner at 20x magnification and on the Leica SP8 confocal at 40x for quantification.

Confocal images were scanned in a z-stack at 1um steps (2um steps for non-neuronal cell types in Fig. S3) and were tiled. The resulting tiling merged image was then processed in FIJI. The maximum intensity projection was taken across 8 planes (5 for FISH of non-neuronal cell types in Fig. S3). Background subtraction was applied with a rolling ball radius of 100 pixels.

## Supplemental Information

**Supplemental Figure 1.**
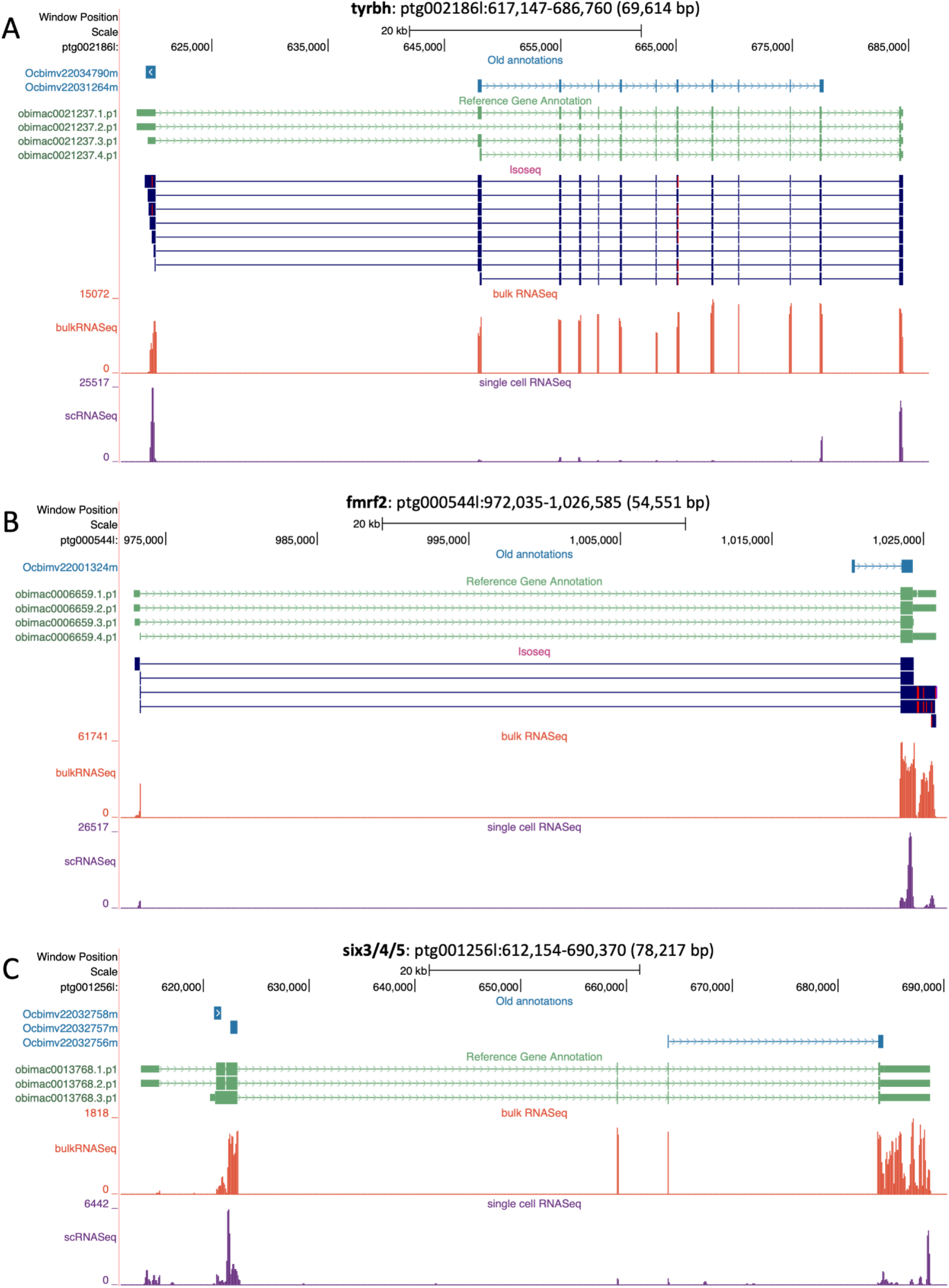
Genome browser output showing improved gene models. Three genes are shown to demonstrate examples of the substantial improvement our gene models achieved with an updated annotation. Each panel contains the following tracks: old gene annotations (blue) from the original *O. bimaculoides* genome annotation all isoforms of new annotations (green), Iso-seq alignment data (dark blue, if present), bulk RNA sequencing alignment data (orange), and single-cell RNA sequencing alignment data (purple). Our single-cell data is poly-A captured, meaning most reads align to the 3’ ends of genes or areas of the genome. (A) **tyrbh**. New gene models (labeled in green, with each potential isoform represented by the obimac gene identifier followed by a period and numeric) include two additional exons compared to the original annotation, one on each end of the gene. These new exons are supported by both Iso-Seq data and bulk RNA-seq data. By including both of these exons, we have captured two piles of scRNA-seq read alignments on each end of the gene. (B) **fmrf2**. New models lengthened the 3’ ends and included an additional exon on the 5’ end compared to the original model. Both of these changes to this gene are supported by Iso-Seq bulk RNA-seq read alignments and improved the amount of scRNA-seq data we are able to capture. (C) **six3/4/5**. This gene was not present in our Iso-Seq data, but shows an example of where the new gene model substantially improved annotation of this region. The new models lengthened the 3’ end and added two exons in the 5’ direction, allowing us to capture more of the scRNA-seq data. We were able to stitch together three genes from the old model (Ocbimv22032756m, Ocbimv22032757m, and Ocbimv22032758m) into a single gene model (obimac0013768). If we had used the old gene model for our single cell analysis, then it would appear that three separate genes have similar expression patterns.

**Supplemental Figure 2.**
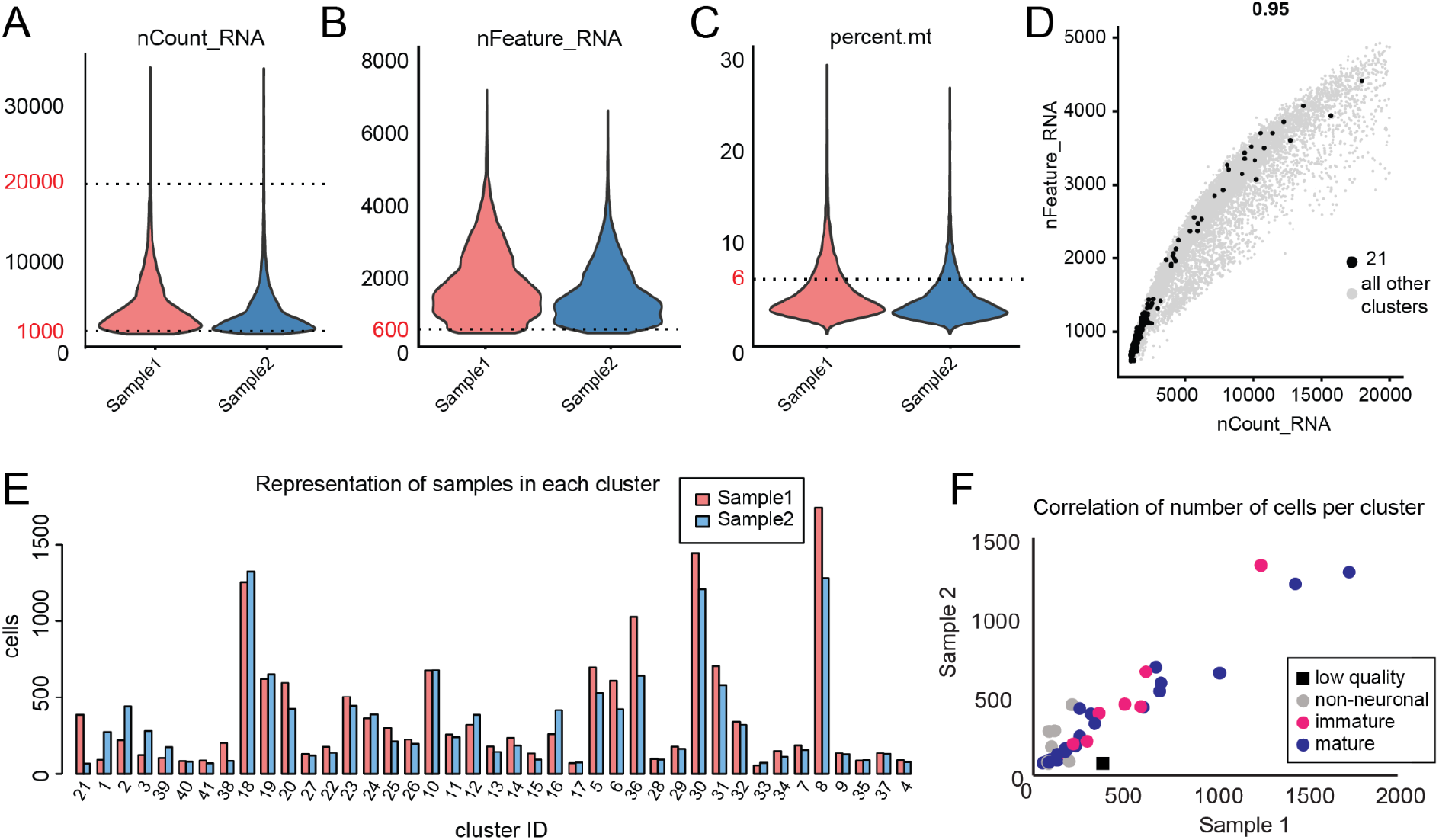
Single-cell RNA sequencing quality control metrics. (A-C) Violin plot showing raw reads (nCount_RNA), genes (nFeature_RNA), and mitochondrial percentage (percent.mt) across the two biological replicates used in this study. Dashed lines and red text indicate thresholds set for downstream processing. (D) Scatter plot showing the correlation between the number of reads and the number of genes for all cells. One cluster (21) had low reads and gene counts and did not have a specific gene expression profile (data not shown). Cluster 21 was therefore excluded from further analysis. (E) Number of cells in each cluster for the two replicates. (F) Scatter plot of number of cells per cluster, with each biological replicate plotted on each axis. Points are color-coded based on whether they represent clusters from non-neuronal, immature, or mature neuronal populations. Neuronal cell types were represented similarly in the two replicates. On the other hand non-neuronal clusters were more variable, perhaps due to their presence in neighboring tissue. The “low quality” cluster (21), which was discarded from analysis, is represented by a black square and was only present in one replicate.

**Supplemental Figure 3.**
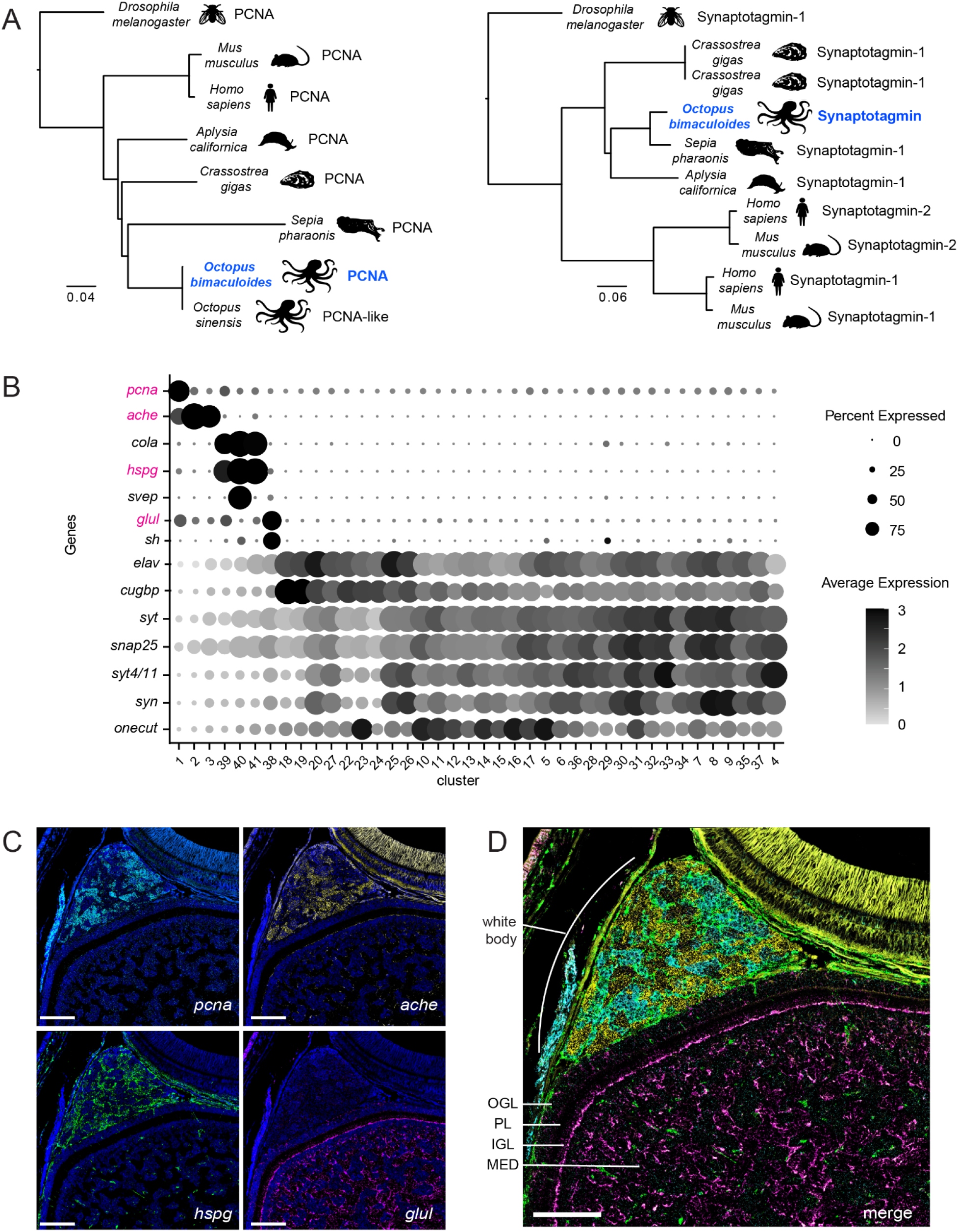
Characterization of non-neuronal clusters. (A) Phylogenetic trees from Orthofinder, demonstrating the method for assignment of gene identities in our updated gene model. (B) Dot plot of conserved markers, showing demarcation of neuronal vs. non-neuronal cells, as well as subtypes within non-neuronal cells. One subset (clusters 1-3) of non-neuronal cells has relatively high expression of markers relating to proliferation and blood (proliferating cell nuclear antigen (*pcna*) and acetylcholinesterase (*ache*)), whereas a second subset (clusters 39-41) are marked by relatively high expression of genes relating to endothelium (collagen type 1 alpha 1 (*cola*), heparan sulfate proteoglycan 2 (*hspg)*, and sushi von willebrand factor type a (*svep*)). Finally, a third cluster (38) has high expression of glutamine synthase (*glul*), which is expressed in glia, and serine hydrolase (*sh*). Markers for these clusters are contrasted with neuronal markers/clusters below. (C) FISH of genes to demonstrate anatomical locations of non-neuronal cell types. *pcna* (top left) marks dividing cells and is localized to the white body, along with *ache* (top right), which is a marker of blood cells. Together, *pcna* and *ache* expression is consistent with the suggested role of the white body in hematopoiesis. *hspg* (bottom left) demarcates endothelial cells and is prominent in the white body as well as the optic lobe, where their morphology resembles vasculature. *glul* (bottom right) is a marker of glia and can be seen within the optic lobe, prominently labeling nuclei on the superficial border of IGL but also extending throughout the neuropil of the medulla. Scale bars indicate 100 um. Nuclei are stained in DAPI. (D) A quadruple FISH of the genes shown in C without nuclei staining. OGL, outer granular layer; PL, plexiform layer; IGL, inner granular layer; MED, medulla. Scale bar indicates 100 um.

**Supplemental Table 1.**
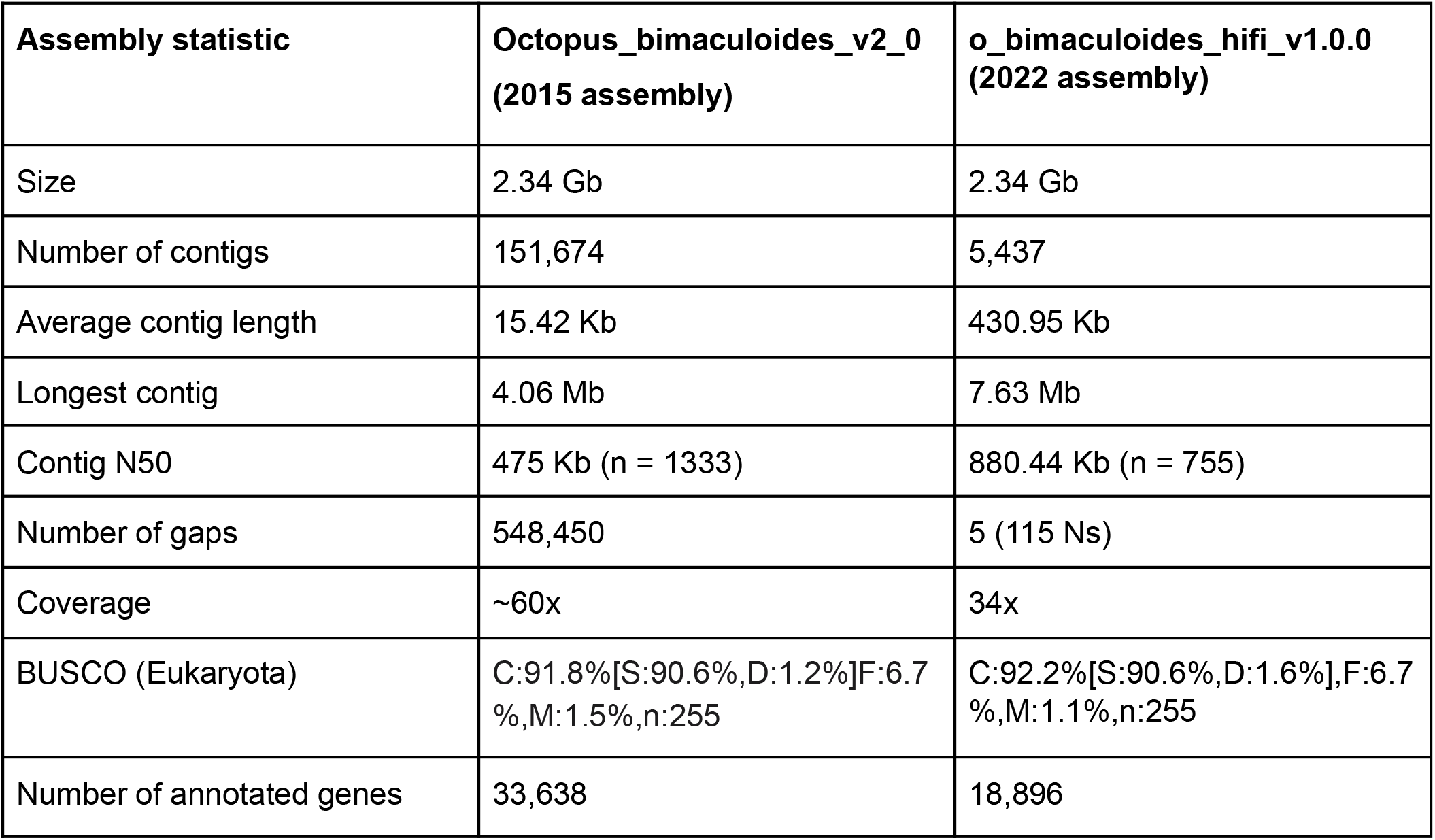
Genome assembly statistics for o_bimaculoides_hifi_v1.0.0 and Octopus_bimaculoides_v2_0. The computational pipeline that led to these results is described in Methods. Benchmarking Universal Single-Copy Orthologs (BUSCO v. 3)^93^ was run against the eukaryota_odb9 database to show an overall completeness of 92.2% (C: complete, S: complete and single-copy, D: complete and duplicated, F: fragmented, M: missing). We used the Eukaryote database rather than Mollusc or Metazoa because those databases contain the original *O. bimaculoides* proteome, thus biasing results. The older gene model contains many fragmented gene annotations, which led to an inflated number of total genes in the original gene annotation file. The new genome annotation contains fewer genes because many of the fragmented annotations have been connected.

| Assembly statistic | Octopus_bimaculoides_v2_0<br>(2015 assembly) | o_bimaculoides_hifi_v1.0.0<br>(2022 assembly) |
| --- | --- | --- |
| Size | 2.34 Gb | 2.34 Gb |
| Number of contigs | 151,674 | 5,437 |
| Average contig length | 15.42 Kb | 430.95 Kb |
| Longest contig | 4.06 Mb | 7.63 Mb |
| Contig N50 | 475 Kb (n = 1333) | 880.44 Kb (n = 755) |
| Number of gaps | 548,450 | 5 (115 Ns) |
| Coverage | ~60x | 34x |
| BUSCO (Eukaryota) | C:91.8%[S:90.6%,D:1.2%]F:6.7<br>%,M:1.5%,n:255 | C:92.2%[S:90.6%,D:1.6%],F:6.7<br>%,M:1.1%,n:255 |
| Number of annotated genes | 33,638 | 18,896 |

**Supplemental Table 2.**
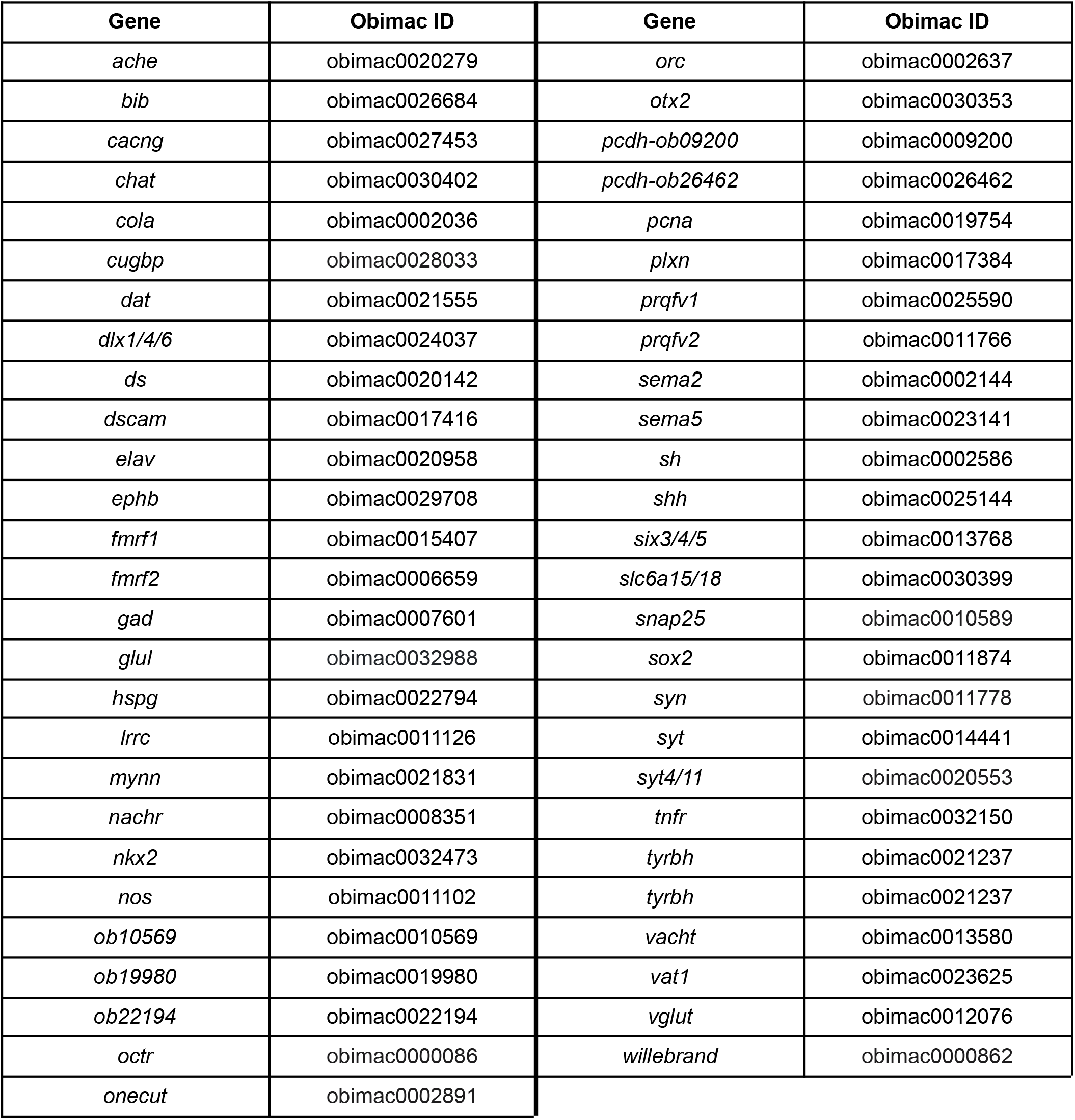
Reference gene table. Obimac gene identifiers and corresponding gene assignment for all genes described in this study. See Methods for more information about gene assignments.

## Supplemental Text. Elucidating unidentified genes

In a number of cases, genes of interest were not well annotated using orthogroup assignment or simple BLAST searches to the non-redundant database. Here we describe additional steps we took to try to assign likely functions to our candidates. In most of these cases orthologous protein predictions could be retrieved from other cephalopod genomes, but not beyond, suggesting that these uncharacterized genes are novel and potentially specific to cephalopods or octopuses. With these in hand, we sought to predict function via deep homology searching as well as structure and function prediction. Homology search and function prediction included the use of the hidden Markov model based tools HHblits and HHpreds^94,95^, and PSI-BLAST^96^. In each case we predicted protein structures using ColabFold^97^, which itself is based on AlphaFold2.0^98^. With these predicted structures in hand, we then searched for distant structural homologues using foldseek^99^.

### obimac0010569

Obimac0010569 has an excellent blast hit to an uncharacterized protein found in *O. vulgaris* and *O. sinensis* called LOC115219258. This hit however combines two separate predicted genes from the previous *O. bimaculoides* genome: obimac0010569 and obimac0010570. While we have iso-seq reads that cover obimac0010570, we have none that link obimac0010569 and obimac0010570. Assuming our gene model has split this locus in error, we created multiple species alignments of the candidate and used the resulting multiple species alignment (MSA) for homology search. HHblits revealed a strong signal of a conserved m13 domain on c-terminal side of the predicted protein, with alignsto proteins like putative m13 family peptidases (e-vals ∼ 1e-100) and endothelin converting enzymes (e-vals ∼ 1e-90). HHPred showed similar signals, with peptidases and in particular metallopeptidases showing homology on the c-terminal end. Interestingly there is some weak homology to Protocadherin-15, but e-values were all >> 0.01

### obimac0022194

We were able to find obimac0022194 orthologs from *O. sinensis, O. vulgaris*, and *O. minor*. Using a multiple sequence alignment from these orthologs as input to HHblits revealed hits to uncharacterized proteins from the cuttlefish, *S. pharaonis* (e-val ∼ 2.2e-15), the limpet *Lottia gigantea* (e-val ∼ 3.9e-8), and slightly weaker hits to a protein annotated as luquin1 in the land slug *Deroceras reticulatum* (e-val 9.9e-8) and a protein annotated as luquin neuropeptide fragment in the annelid polychaete *Platynereis dumerilii* (e-val 2.6e-5). HHpred gave no further insight. We then used AlphaFold2.0 to computationally predict a structure using our MSA. The resulting structures were generally low confidence and largely disordered. A representative view of the rank1 model is shown below in Fig. ST1. We further searched for structural homologs using foldseek, but zero hits were returned.

**Figure ST1.**
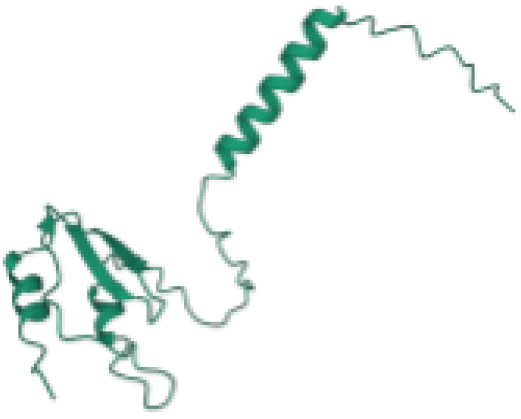
AlphaFold prediction for obimac0022194 showing the structure of the rank1 model.

### obimac0019980

Using protein BLAST searches against individual cephalopod genomes we were able to find an *O. sinensis* ortholog to obimac0019980 with strong homology (e-val ∼ 3e-88) and only a very weak hit to the cuttlefish *Sepia officinalis* (e-val = 0.003). We used pairwise alignments of the *O. bimaculoides* and *O. sinensis* proteins as input to HHblits and HHpred. HHblits recovered no significant homology. HHpred similarly did not recover orthologs in distant genomes but did predict a transmembrane domain and a signal peptide sequence within obimac00119980. We used DeepTMHMM^100^ to confirm the presence of the signal peptide and transmembrane domain. A clear signal peptide is present and DeepTMHMM classifies this protein as globular + signal peptide (Fig. ST2). We further used AlphaFold2.0 to provide a predicted structure. This structure was largely disordered but two clear helical regions are revealed. Topology searching using foldseek revealed no similar structures.

**Figure ST2.**
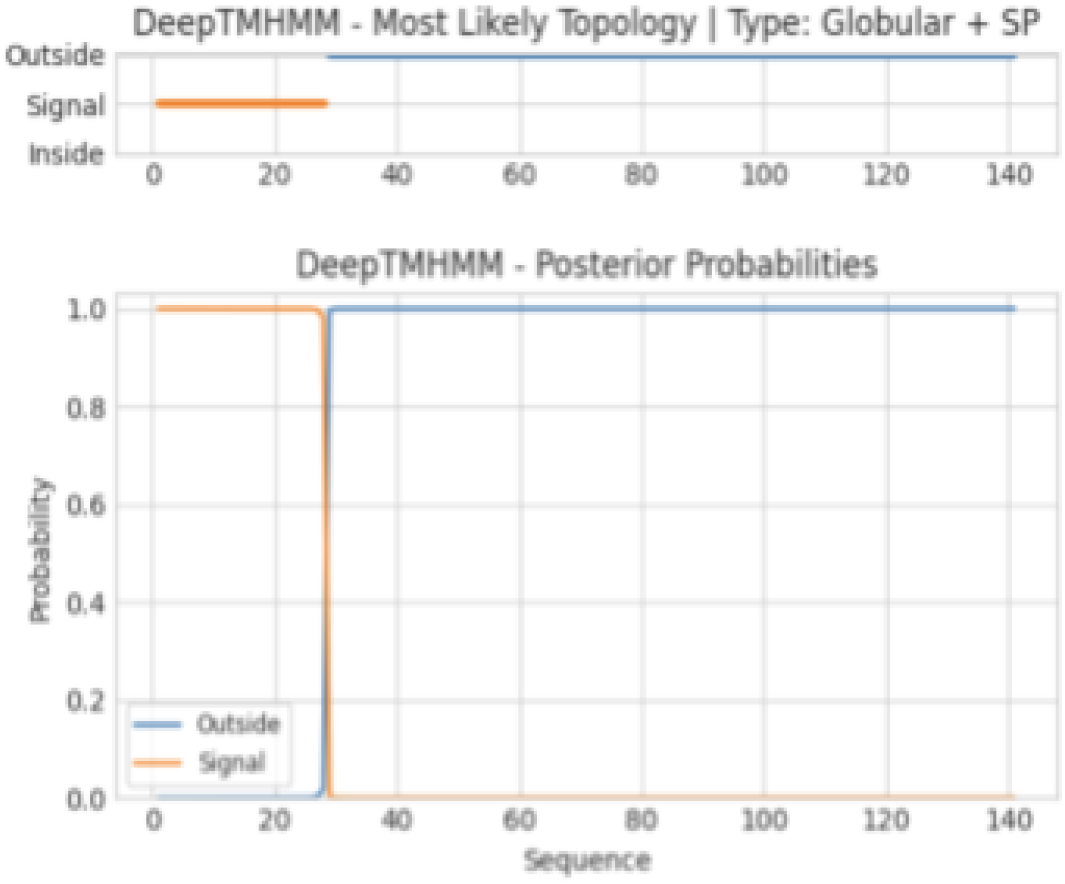
Evidence of a globular protein plus signal peptide structure for obimac0019980 as predicted by DeepTMHMM^100^.

